# Post-transcriptional regulatory feedback encodes JAK-STAT signal memory of interferon stimulation

**DOI:** 10.1101/2022.05.13.489832

**Authors:** Eirini Kalliara, Malgorzata Kardyńska, James Bagnall, David G. Spiller, Werner Muller, Dominik Ruckerl, Subhra K Biswas, Jarosław Śmieja, Pawel Paszek

**Author notes:** **Correspondence:** Pawel Paszek. These authors contributed equally (Eirini Kalliara, Malgorzata Kardyńska).

## Abstract

Immune cells fine tune their responses to infection and inflammatory cues. Here, using live-cell confocal microscopy and mathematical modelling, we investigate interferon induced JAK-STAT signalling in innate immune macrophages. We demonstrate that transient exposure to IFN-γ stimulation induces a long-term desensitisation of STAT1 signalling and gene expression responses, revealing a dose- and time-dependent regulatory feedback that controls JAK-STAT responses upon re-exposure to stimulus. We show that IFN-α/β1 elicit different level of desensitisation from IFN-γ, where cells refractory to IFN-α/β1 are sensitive to IFN-γ, but not *vice versa*. We experimentally demonstrate that the underlying feedback mechanism involves regulation of STAT1 phosphorylation but is independent of new mRNA synthesis and cognate receptor expression. A new feedback model of the protein tyrosine phosphatase activity recapitulates experimental data and demonstrates JAK-STAT network’s ability to decode relative changes of dose, timing, and type of temporal interferon stimulation. These findings reveal that STAT desensitisation renders cells with signalling memory of type I and II interferon stimulation, which in the future may improve administration of interferon therapy.

## Introduction

Immune signalling systems decode external signals in order to produce appropriate responses (Levine, Lin, and Elowitz 2013). Underlying mechanisms involve complex regulatory networks with feedback loops that generate outputs depending on the signal type, strength, or duration (Bhalla and Iyengar 1999; Novak and Tyson 2008). For example, the temporal responses of the nuclear factor -κB (NF-κB) transcription factor encode different pathogen-derived molecules and cytokines of the immune system (Adelaja et al. 2021; Martin et al. 2020; Kellogg et al. 2017) and adapts to rapidly changing inflammatory cues (Adamson et al. 2016; DeFelice et al. 2019; Ashall et al. 2009; Tay et al. 2010; Son et al. 2021). The emergent properties of the cellular signalling networks serve as a paradigm to understand how immune cells process inflammatory cues (Dorrington and Fraser 2019).

Interferons (IFNs) are secreted signalling molecules with antiviral, antiproliferative and immunomodulatory functions in response to infection (Platanias 2005). IFN-γ is a type II interferon, produced by innate immune natural killer cells and innate lymphoid cells as well as T lymphocytes of the adaptive immunity, mainly, but not only, in the direct response to pathogens (Ivashkiv 2018). Innate immune macrophages are the main physiological target of newly secreted IFN-γ (Epelman, Lavine, and Randolph 2014). IFN-γ exerts its biological function trough binding to its cognate receptor (IFN-γ R), which activates Janus kinase (JAK)-STAT signalling pathway (Villarino, Kanno, and O’Shea 2017). IFN-γ binding leads to the nuclear translocation of Signal Transducers and Regulators of Transcription 1 (STAT) homodimers, which directly activate expression of hundreds of interferon-regulated genes (ISGs) via conserved sequences in their promoters (Barrat, Crow, and Ivashkiv 2019). Functionally related type I interferons, such as IFN-α and IFN-β regulate overlapping sets of genes via ISGs (in part through STAT1 homodimers), but also utilise IFN-stimulated response elements (trough STAT1-STAT2-IRF9 complexes) (Van Eyndhoven, Singh, and Tel 2021). Regulation of IFN signalling exemplifies the intricate balance within the immune system to produce appropriate responses. A lack of IFN-γ responses results in susceptibility to pathogen infection (Dalton et al. 1993; Jouanguy et al. 1997), thus IFN-γ has been clinically used to treat inflammation including sepsis (Payen et al. 2019). In turn, a sustained IFN signature has been associated with autoinflammatory diseases such as arthritis (Ivashkiv 2018) or cancer (Barrat, Crow, and Ivashkiv 2019) and uncontrolled type I IFN responses have also recently been associated with severe COVID 19 symptoms (Blanco-Melo et al. 2020; Lee and Shin 2020). The properties of the JAK-STAT network are controlled via the temporal regulation of STAT activity via the receptor availability (Bach et al. 1995) and regulatory feedback (Villarino, Kanno, and O’Shea 2017). Known feedback mechanisms involve transcriptional activation of Suppressors of Cytokine Signalling (SOCS), protein inhibitor of activated STAT (PIAS) and ubiquitin-specific peptidase (USP18) (Kok et al. 2020; Mudla et al. 2020; Ivashkiv 2018; Yasukawa, Sasaki, and Yoshimura 2000; Liau et al. 2018; Smieja et al. 2008) as well as post-translational regulation of tyrosine phosphatase activity (Böhmer and Friedrich 2014). To achieve the appropriate level of response, innate immune cells acquired the ability to adapt to repeated immune challenges, i.e., ‘memory’ (Netea et al. 2020). IFN-γ sensitises cells for subsequent stimulation through long-term epigenetic changes (Kamada et al. 2018) as well as regulation of STAT expression (Hu et al. 2002). In contrast, pathway desensitisation represents a mechanism that prevents prolonged or uncontrolled activation to persistent stimulation. Best characterised examples involve endotoxin resistance in the toll-like receptor system (Morris, Gilliam, and Li 2014; Buckley, Wang, and Redmond 2006), however desensitisation has also been extensively studied for type I interferon signalling (Mudla et al. 2020; Sakamoto et al. 2004; Kok et al. 2020). How cells adapt to temporal IFN-γ stimulation is less understood.

In this work, we use live-cell confocal microscopy and mathematical modelling to investigate STAT1 responses to IFNs in innate immune macrophages. Using pulses of IFN-γ and IFN- α/β1 at different concentrations and frequency we demonstrate a long-term dose-, time- and stimulus-specific desensitisation of STAT1 signalling and gene expression responses. We demonstrate that pathway desensitisation involves control of STAT1 phosphorylation and is independent of new mRNA synthesis and IFNγ R expression. Our new dynamical mathematical model of the JAK-STAT signalling network that recapitulates our experimental data, demonstrates that stimuli-induced STAT1 desensitisation renders cells with signalling memory of IFN stimulation. These analyses reveal the ability of macrophages to quantitatively fine-tune their responses to temporal interferon stimulation.

## Materials and methods

### Cells and reagents

Immortalised bone marrow-derived mouse macrophage (iBMDM) cells (Hornung et al. 2008) were cultured in Dulbecco’s Modified Eagle Medium (DMEM) with L-glutamine (Sigma-Aldrich), supplemented with 10 % (v/v) of heat-inactivated Foetal Bovine Serum (Life Technologies Ltd) and 1 % (v/v) of Penicillin-Streptomycin (Sigma-Aldrich) solution. Cells were cultured between passages 6-30 in sterile NuncTM 10 cm cell culture petri dishes (ThermoFisher Scientific) till 80-90 % confluent. Lentiviral transduction (Bagnall et al. 2015) was used to develop reporter iBMDM line constitutively expressing murine STAT1 coding sequence C-terminally fused with the red fluorescent protein (STAT1-tagRFP) and murine STAT6 coding sequence N-terminally fused with yellow fluorescent protein (Venus-STAT6). Additionally, cells expressed histone H2B protein fused with cyan fluorescent protein (AmCyan-H2B). iBMDMs were sequentially transduced and triple positive cells were identified using fluorescence single cell sorter (BD Influx) to derive a clonal reporter line. Cells were stimulated with recombinant mouse IFN-γ (575306, Biolegend), IL-4 (574306, Biolegend), IFN-α (752802, Biolegend) and IFN-β1 (581302, Biolegend).

### Confocal microscopy and image analysis

Fluorescent confocal imaging was performed with Zeiss LSM 710, LSM 780 and LSM 880 laser scanning confocal microscopes, which collect emitted signals using dichroic mirrors and band-pass filters or spectral separation and detector arrays. Fluorophores were exited with the appropriate laser lines (AmCyan excited with 458 nm laser line, Venus with 514 nm laser line, tagRFP with 561 nm laser line). Imaging was conducted using Fluar 40x NA 1.3 (oil immersion) objective using Zen 2010b SP1 software. Time-lapse images were performed by seeding cells onto 1-compartment or 4-compartment round glass-bottom 35 mm culture dishes (627860 & 627870, Greiner Bio-One) at density of 300 × 10^3^ cells/dish or 100 × 10^3^ cells/compartment, respectively. Imaging plates were placed on the microscope stage in a humidified incubator maintaining 37 °C and 5% CO_2_. Image series were captured with a time interval of 5 mins by selecting several regions of interest per condition. Image analysis was performed using Imaris Bitlane software version 9.3 using AmCyan fluorescent signal was used as a nuclear mask to segment and track single cells. Nuclear masking was tailored according to nuclei size and cell movement depending on the experimental conditions. Automated cell tracking was executed by Autoregressive Motion Imaris algorithm. Quantified data were extracted as the nuclear mean fluorescence intensity for the respective fluorescent channels under investigation. The values were imported to GraphPad Prism 9 for further processing and statistical analyses.

### Live-cell luminometry

Lentiviral GAS-luciferase reporter (GAS-luc) construct was generated from the previously described 5xκB-Luc plasmid (Brignall et al. 2017). Namely, Pac1 and Xho1 restriction sites were used to replace κB elements with a GAS consensus sequence (AGTTTCATATTACTCTAAATCAGTTTCATATTACTCTAAATCAGTTTCATATTACTCTAAATCAGTTTCATATTACTCTAAATCAGTTTCATATTACTCTAAATC) (Satoh and Tabunoki 2013). Lentivirus production and transduction of iBMDMs were carried out as previously described (Bagnall et al. 2015). For the purposes of live-cell luminometry 10 × 10^3^ GAS-luc cells were seeded onto white, flat-bottom 12-well culture microplates (Greiner Bio-one) in 1 ml complete medium. 10 μl of 100 mM D-luciferin (Biosynth) was added 24 h prior to the start of the assay. Live-cell luminometry was performed using a Fluostar Omega luminometer (BMG Labtech) at 37 °C and 5% CO^2^. Light production was measured at 492 nm with 10 mins intervals for 24 h.

### Single molecule RNA-FISH

Custom smFISH probes were designed using Stellaris Probe Designer version 4.2 (Biosearch Technologies Inc.) against murine STAT1 (NM_001205313) and SOCS1 (NM_001271603) (see Table S1 for the probe sets). Each probe was attached with a fluorophore (either Quasar 570 or Quasar 670). Cells were seeded onto 18 mm coverslips (BDH) placed in 6-well or 12-well plates (Corning, Appleton Woods Limited). For the measurement, cells were washed with PBS and fixed with 4 % formaldehyde (Sigma-Aldrich) for 10 mins. Subsequently, cells were permeabilised with 70% Ethanol (EtOH) and left for at least 1 h at 4 °C according to Stellaris protocol for adherent cells (Orjalo and Johansson 2016). Probe hybridisation was performed at a concentration of 125 nM for up to 16 h at 37 °C in a humidified chamber. Coverslips were mounted on microscope slides (ThermoFisher) using Vectashield mounting medium (Vector Laboratories) containing 4’, 6-diaminidino-2-phenylinode (DAPI) for nuclei staining. Imaging was performed using Deltavision deconvolution system equipped with a Plan Apo 60x 1.42 NA (oil immersion) objective. Light-emitting diodes were used to illuminate specimens with the desired excitation wavelength band (358 nm for DAPI, 570 nm for Quasar 570 and 670 nm for Quasar 670). Images were acquired as z-series with an optical spacing of 0.2 μm using MetaMorph acquisition software, respectively. Obtained images were deconvolved using Huygens Professional software. mRNA quantification was performed in FISHQuant (Tsanov et al. 2016).

### qRT-PCR

Cells were seeded onto 6-well plates with a density of 300 × 10^3^ cells/well. RNA extraction was performed using Qiagen RNeasy mini kit according to manufacturer instructions. RNA concentration was quantified using a NanoDrop 1000 Spectrophotometer (Thermo Fisher Scientific). 1 μg of RNA was used for reverse transcription using the SensiFAST cDNA synthesis kit (Bioline). qPCR reactions were prepared in MicroAmp Fast Optical 96-well plates with barcode (Applied Biosystems) using SYBR Green master mix (Applied Biosystems). A 10 μl final volume reaction was prepared, which included 5 μl of the SYBR Green master mix, 0.5 μl of primer, 0.5 μl of the cDNA template and 3.5 μl of sterile DNase-free water. Amplification was performed using the StepOne Plus Real-Time thermal cycler (Applied Biosystems). Amplification of β-actin gene was used as a control across all samples. Genes of interest were first normalised to β-actin expression within the same sample. Normalised expression of target genes in the samples of interest were then compared to the normalised expression of the same genes in a reference control sample (untreated cells). Final levels of gene expression are presented as a fold change in the target sample compared to the reference/control sample (ΔΔCt) following established methodology (Livak and Schmittgen 2001). All samples were examined in three technical replicates. The following primer sequences were used (5’ to 3’): β-Actin TATCCACCTTCCAGCAGATGT (forward) and AGCTCAGTAACAGTCCGCCTA (reverse); STAT1 TCATCCCGCAGAGAGAAC (forward) and TGAAACGACCTAGAAGTGAG (reverse); PD-L1 GAAAGTCAATGCCCCATACC (forward) and ATTGAGAAGCATCCCCTCTG (reverse), SOCS1 GAGACCTCATCCCACCTCTC (forward) and AGACACAAGCTGCTACAACC (reverse), CXCL10 CACGTGTTGAGATCATTGCC (forward) and TCACTCCAGTTAAGGAGCCC (reverse), ARG1 CTGTCTTTTAGGGTTACGGC (forward) and CTCGAGGCTGTCCTTTTGAG (reverse) and TNFA, TGAGGTCAATCTGCCCAAGT (forward) and TGGACCCTGAGCCATAATCC (reverse)

### Immunoblotting

Cells were lysed in 80 μl of radioimmunoprecipitation assay buffer (RIPA) (89900, ThermoFisher, Scientific) supplemented with Pierce Protease Inhibitor Mini tablets (A32953, ThermoFisher Scientific) as per manufacturer’s instructions. Cell extracts were incubated for 15 mins on ice followed by centrifugation at 12000 rpm for 12-15 mins at 4 °C. Protein concentration of each sample was measured with Pierce Bicinchoninic acid (BCA) protein assay kit (23227, ThermoFisher Scientific) according to manufacturer’s protocol. Polyacrylamide gel of 10 % size pore was prepared using a 30 % w/v Acrylamide stock solution (A2-0084, Geneflow). A 2x protein loading buffer was freshly prepared using 950 μl of 2x Laemmli buffer (161-0737, Bio-Rad) mixed with 50 μl of β-ME. 20 μg of protein sample was mixed with the appropriate amount of protein loading buffer and sterile double distilled water (ddH2O). Diluted protein samples were denatured at 95 °C for 5 mins and loaded onto the wells of polyacrylamide gel. 5 μl of Precision Plus Protein Dual Colour (1610374, Bio-Rad) ladder was also loaded onto the gel and run alongside the samples to determine the molecular weight. Electrophoresis was performed at 120 V for 60-90 mins. Transfer of proteins was confirmed by staining of membranes with Ponceau S stain (0.1 % w/v Ponceau S solution in 1% v/v Acetic acid). Membranes were washed in PBS-Tween 20 (PBS-T) for 5 mins three times and blocked in 5 % non-fat powdered milk (Sigma-Aldrich) in PBS for 1 h at room temperature. Probing with primary antibodies was conducted o/v at 4 °C. The next day, membranes were washed in PBS-T for 5 mins three times to remove unbound primary antibody and subsequently blocked in HRP-conjugated secondary IgG antibody for 1 h at room temperature. Following incubation, excess of secondary antibody was removed by washing membranes in PBS-T for 5 mins three times. Pierce ECL Western Blotting Substrate (32106, ThermoFisher Scientific) was used to incubate membranes as per manufacturer’s instructions. Luminescent signal was captured on Carestream Biomax Xar films (F5513, Sigma-Aldrich) using an automatic X-ray processor model JP-33 (JPI). Primary antibodies were purchased from Cell Signaling Technology®: STAT1 rabbit polyclonal antibody (9172S) used at 1:1000 dilution; phospho-STAT1 (pTyr701) (58D6 clone) rabbit monoclonal antibody (9167S) at 1:1000 dilution; and β-actin (13E5 clone) rabbit monoclonal antibody (4970S) at 1:2000 dilution. Primary antibodies were detected with Horseradish Peroxidase (HRP)-conjugated goat anti-rabbit IgG (H+L) secondary antibody (65-6120, ThermoFisher Scientific) using 1:3000 dilution. Quantification of immunoblotting was performed in ImageJ software using the ‘Measure’ function with setting ‘Mean gray value‘ applied to individual bands (relative to the β-actin loading control).

### FACS analysis of receptor expression

Cells were either untreated (control) or treated with continuous 100 ng/ml IFN-γ, or 1 h pulse of 100 ng/ml of IFN-γ or combined pulse of IFN-α/β1 (50 ng/ml of each). After scraping, collected cell suspensions were centrifuged at 400 x g for 5 mins and resuspended in 5 ml of PBS. For each condition, 1 × 10^6^ cells were transferred in FACs tubes (STEMCELL Technologies). First, cells were stained with viability dye (Zombie Aqua™ fixable viability dye, Biolegend) in 1:1000 dilution in PBS and they were incubated at room temperature, in the dark, for 15 mins. Cells were then washed with MACs buffer (0.5% BSA and 250 μM EDTA in PBS) and centrifuged at 400 x g for 5 mins. Next, cells were stained with PE-conjugated primary antibodies against IFNγ-R α chain (clone GIR-208, Biolegend) and ΙFNγ-R β chain (clone 2HUB-159, Biolegend) at a final concentration of 5 μg/ml in MACs buffer. Incubation was performed at room temperature, in the dark, for 25-30 mins. Stained cells were then washed with MACs buffer and centrifuged at 400 x g for 5 mins. Unstained cells served as control. Subsequently, cells were fixed using 4 % of paraformaldehyde (PFA) solution for 10 mins and washed with PBS and centrifuged at 400 x g for 5 mins. Finally, cells were resuspended in 500 μl of PBS and kept at 4 °C o/n. Samples were analysed the next day using a BD Fortessa X20 flow cytometer. PE-conjugated IFNγ-R α and β antibodies were excited using a blue (488 nm) laser line, while viability dye was excited with a violet (405 nm) laser line. Data were analysed using FlowJo (version 10.3.0) and statistical analysis was performed using GraphPad Prism 8 (version 8.4.2). Data are presented as Geometric mean of fluorescence intensity acquired from three repeated experiments.

### JAK-STAT model development

The JAK-STAT mathematical model incorporating type I and type II interferon signalling was developed based on the existing model of IFN-β induced signalling pathway (Smieja et al. 2008). The original model consisted of 27 ordinary differential equations (ODES), but some molecules which were not relevant for the current work were removed, these included action of hypothetical protein Phy as well as TAP1 and LMP2 mRNA output. The model was subsequently expanded to incorporate additional variables and interactions, including extracellular IFN-α/β1 and ΙFN-γ, their cognate receptors as well as the activation and the action of PTP proteins, including formation of PTP complexes with STAT1. Three auxiliary variables were created to model PTP activation, resulting in total of 37 ODEs. Model parameters were then fitted to recapitulate experimental data including all the time-lapse microscopy for IFN-γ and IFN-α/β1 stimulation as well as levels of STAT1 mRNA and protein (see Tables S2 to S5 for model equations, parameters and initial conditions). Simulations were performed using MATLAB R2020b. To directly compare the experimental results with model simulations, nuclear STAT1-tagRFP trajectories were scaled from arbitrary fluorescence levels to number of molecules. First, baseline florescent levels were removed (by subtracting the minimum value in the dataset). Resulting levels were subsequently multiplied by a scaling factor to match the maximal level of nuclear STAT1 in simulations, across different experimental protocols.

### Statistical analyses

Statistical analysis was performed using GraphPad Prism 8 software (version 8.4.2). The D’Agostino-Pearson test was applied to test for normal (Gaussian) distribution of acquired data. Two-sample comparison was conducted using non-parametric Mann Whitney test, for analyses of variance Kruskal-Wallis ANOVA with Dunn’s multiple comparisons test was performed. Non-parametric Spearman correlation was conducted to test association between two selected variables coefficient of correlation r.

## Results

### IFN-γ induces desensitisation of STAT1 signalling and gene expression responses

To investigate STAT signalling, we engineered a reporter murine immortalised bone marrow-derived macrophage (iBMDM) line (Hornung et al. 2008) constitutively expressing STAT1 fused to red fluorescent protein (STAT1-tagRFP). Reporter cells also expressed the nuclear marker AmCyan-H2B to enable automated segmentation and tracking of confocal microscopy images. In addition, as a tool to study macrophage activation, reporter cells expressed STAT6 tagged with yellow fluorescent protein (Venus-STAT6). We focused on interferon signalling and assayed STAT1 activation via confocal microscopy. Untreated cells exhibited a predominant cytoplasmic localisation of STAT1-tagRFP, but a continuous simulation with a saturated dose of 100 ng/ml of IFN-γ (see Fig. S1A for a dose response) resulted in a single transient nuclear STAT1 translocation (Fig. 1A and B and Movie 1). Cells exhibited a maximal STAT1-tag RFP nuclear localisation at 72 ±42 mins (mean ± standard deviation, SD) after the start of the experiment (Fig. 1C). The translocation lasted for up to 6 h, after which nuclear STAT1-tagFRP levels returned towards the pre-stimulation steady-state. A transient activation in response to continuous treatment is a hallmark of desensitisation, where cells become unresponsive to prolonged presence of stimulus (Morris, Gilliam, and Li 2014; Buckley, Wang, and Redmond 2006). In contrast, other signalling systems remain active if the stimulus is present, as in the case of the cytokine stimulation of the NF-κB system (Nelson et al. 2004; Hoffmann et al. 2002; Ashall et al. 2009). To understand whether STAT1 signalling exhibited desensitization, we treated cells with a single 1 h pulse of 100 ng/ml of IFN-γ. We found that STAT1-tagRFP translocation kinetics were similar to that of the continuous treatment (Fig. 1A, B and C). There were no differences in peak amplitude of the nuclear STAT1-tagRFP, while differences in the AUC and peak timing were <15% (Fig. 1C). To quantify the level of desensitisation following a single IFN-γ pulse, a second 1 h pulse of 100 ng/ml of IFN-γ was applied 6 hours after the end of the first pulse (referred here as 6 h pulsing interval). We found that cells were refractory to the second IFN-γ pulse as they exhibited no detectable STAT1 activation (Fig. 1A-C). The lack of STAT1-tagRFP activation was also observed when the pulsing interval was extended to up to 24 h, either in iBMDMs assayed on the microscope for the entire duration of the experiment or only during the second (and third pulse) (Fig. 1D).

**Figure 1.**
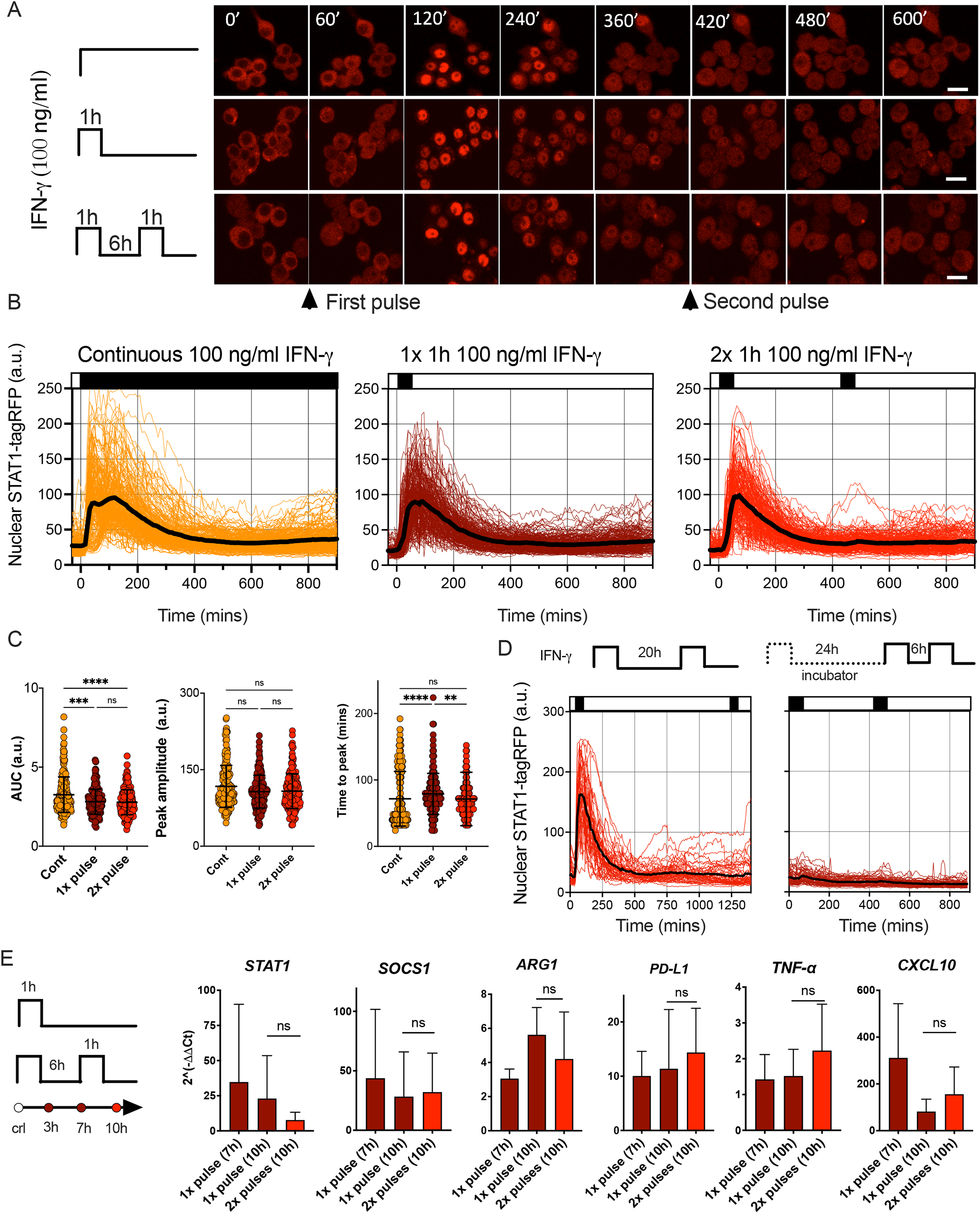
IFN-γ induces desensitisation of STAT1 signalling. A. Representative confocal microscopy images of STAT1-tagRFP iBMDMs cells stimulated with continuous, 1h pulse or two 1h pulses of 100ng/ml of IFN-γ at 6h interval. Time stamp in mines, scale bar 10 μm. B. Temporal STAT1-tagRFP trajectories in reporter iBMDMs in response to different IFN-γ treatment protocols (as indicated on the graph). Shown are individual nuclear STAT1-tagRFP trajectories (colour-coded according to treatment protocol) as well as ensemble average (in black) for 224, 216 and 204 cells for continuous, 1h pulse or two 1h pulses at 6h interval, treatment protocol, from three replicates, respectively. STAT1-tagRFP fluorescence shown in arbitrary units (a.u.), time in minutes (mins). C. Characteristics of single cell nuclear STAT1 trajectories presented in B. From the left: distributions of AUC (over 10h), peak amplitude, and time to peak under different treatment conditions. Individual cell data depicted with circles (with mean ± SD per condition) and colour-coded according to treatment protocol. Kruskal-Wallis one-way ANOVA with Dunn’s multiple comparisons test was used to assess differences between groups (*p < 0.05, **p < 0.01, ***p < 0.001, ****p < 0.0001, ns – not significant). D. Temporal STAT1-tagRFP trajectories in reporter iBMDMs in response to long interval IFN-γ pulsing protocols. Shown are individual nuclear STAT1-tagRFP trajectories as well as ensemble average (in black) for 49 and 67 cells treated with two 1h 100 ng/lm IFN-γ pulses at 20h interval (an imaged under the microscope throughout the experiment) or two pulses at 24h interval (and additional pulse 6h later) while maintaining cells in the incubator before the second pulse. STAT1-tagRFP fluorescence shown in arbitrary units (a.u.), time in minutes (mins). Data from two replicates. E. Fold change of STAT1, SOCS1, ARG1, PD-L1, TNF-α and CXCL10 gene expression response as assessed by qRT-PCR. Wild type iBMDMs stimulated with a 1h pulse of 100ng/ml IFN-γ, or two 1h pulses of 100ng/ml IFN-γ at 6h interval. Shown is the mean fold change (2^(-ΔΔCt), against unstimulated controls) and SD from three replicates measured at 7 and 10 h after the start of the experiment, respectively. Non-parametric Mann Whitney test was used to determine statistical significance between conditions (ns-not significant, p>0.05).

Imaging approaches provide insights into response variability. While all cells responded to saturating IFN-γ treatment, STAT1-tagRFP trajectories exhibited cell-to-cell variability in terms of the AUC and peak amplitude (Fig. 1C). Notably, both showed significant positive correlations with resting nuclear or cytoplasmic STAT1 levels (at t=0 mins) (Fig. S1B and C). We also found significant correlations between the AUC of the response to 1^st^ vs. 2^nd^ pulse (r=0.6, p-val<0.0001) and AUC of the response to the 2^nd^ pulse vs. the nuclear STAT1-tagRFP levels before treatment (at 420 mins, Fig. S1C). This demonstrates that the level of the stimuli-induced STAT1 activation is proportional to the total (and nuclear) resting levels, while the observed variability is likely associated with cell intrinsic differences in STAT1 expression between cells, as demonstrated for other signalling systems (Lee et al. 2014; Kardynska et al. 2018; Patel et al. 2021). This is consistent with recent analyses of IFN-γ signalling demonstrating that phenotypic variability rather than random noise controls heterogeneity of STAT1 responses (Topolewski et al. 2022).

IFN-γ regulates hundreds of target genes, which rely on STAT1-dependent transcription (Barrat, Crow, and Ivashkiv 2019). To evaluate whether desensitization of STAT1 signalling resulted in functional inhibition of inducible gene expression, population-level qRT-PCR was performed (Fig. 1E). We found that IFN-γ upregulated expression across a panel of 6 genes (at 7 h and 10 h following the 1 h pulse of 100 ng/ml of IFN-γ). Consistent with desensitisation, the expression of *Stat1, Socs1*, and *Arg1* mRNA following the second 1 h pulse of 100 ng/ml of IFN-γ at the 6 h interval showed no further induction in comparison to cells stimulated with a single pulse, measured at 10 h from the beginning of the experiment. A subset of genes, namely *Cxcl10, Tnf-α* and *Cd274*, exhibited some but limited (not statistically significant) induction in response to the second pulse, which might reflect their multimodal transcriptional regulation (Falvo, Tsytsykova, and Goldfeld 2010; Vazirinejad et al. 2014; Liu et al. 2000). Overall, these analyses demonstrate that a pulse IFN-γ stimulation induces a long-lasting desensitisation of STAT1 signalling and gene expression responses, which renders cells refractory to stimulus upon re-exposure.

### Desensitisation of STAT1 signalling depends on dose and timing of IFN-γ

To further investigate the regulation of desensitisation, reporter iBMDMs were treated with two pulses of IFN-γ at 6 h intervals, such that the concentration of the first pulse was varied (100, 10, 5 and 1 ng/ml), while the concentration of the second pulse was constant at 100 ng/ml (Fig. 2A). In response to the varied concentration of the first pulse, STAT1-tagRFP activity showed a dose-dependency, with each consecutive nuclear peak amplitude (P1) being significantly higher as the dose increased (Fig. 2B). The response to the second 100 ng/ml pulse was also varied, but we found that it was determined by the amplitude of the first response, i.e., the higher the initial response the lower the response to the 2^nd^ pulse. This was marked by significant increases in the peak nuclear STAT1-tagRFP amplitude (P2) as the dose of the first pulse decreased (except for the 100-100 pulsing regime where only few cells exhibited detectable responses to the second pulse). The responses to the first and second pulse measured as peak nuclear STAT1-tagRFP showed significant positive correlations across doses (of at least r=0.4, p<0.0001, Fig. 2B), suggesting an intrinsic ability of some cells to respond more robustly to both pulses. We found that the saturated 100 ng/ml dose (in the first pulse) produced a significantly higher nuclear STAT1-tagRFP response in terms of the AUC, when compared to other pulsing protocols over the 840 mins duration of the experiment (Fig. 2C). However, when stimulated with non-saturating doses (in the first pulse), cells exhibit the same AUC irrespectively of the concentration of the first pulse. This suggest that the overall temporal STAT1 response to multiple IFN-γ inputs is inherently restricted.

**Figure 2.**
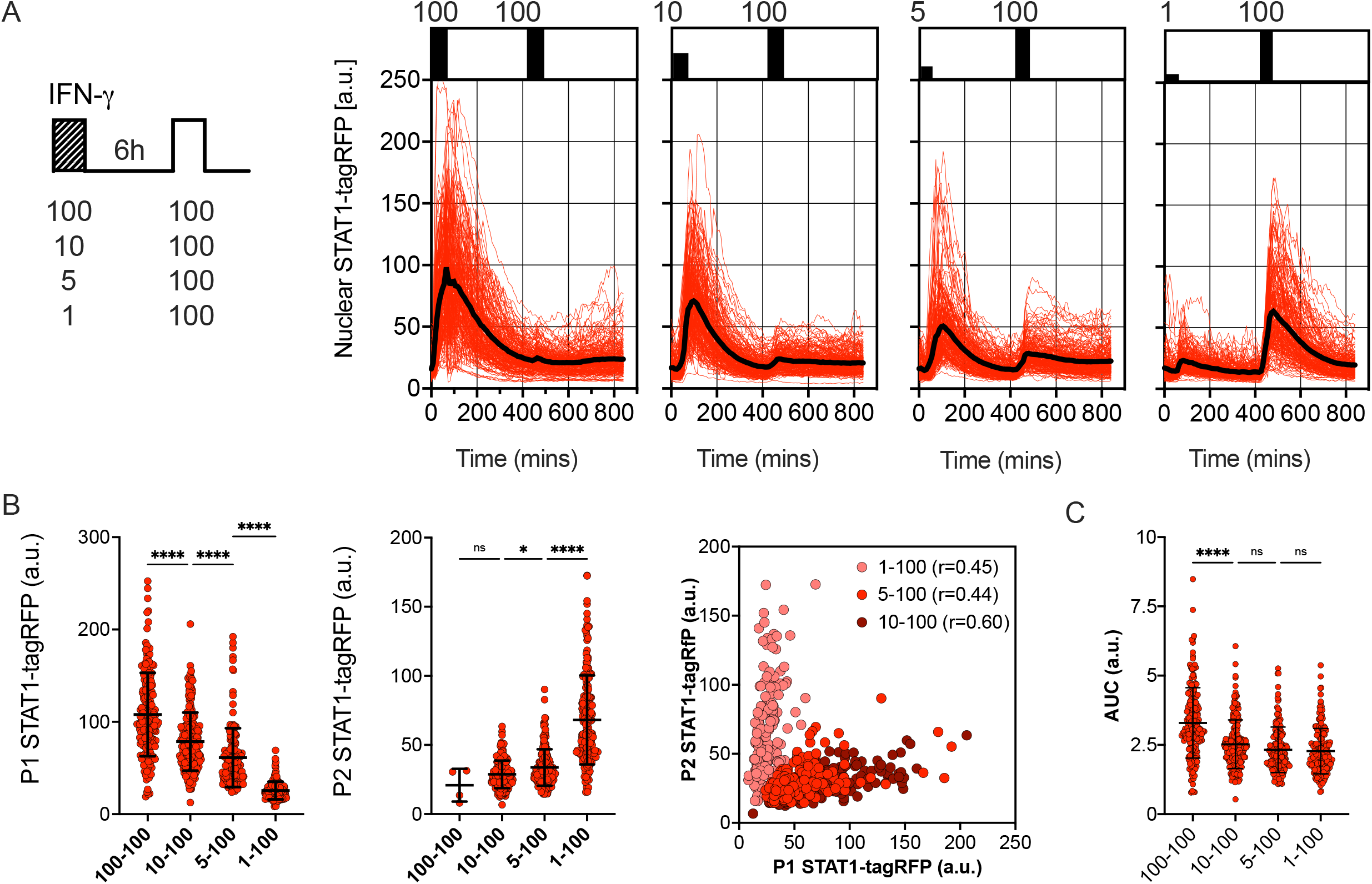
Desensitisation of STAT1 signalling is dose-dependent. A. Single cell analyses of STAT1 responses to IFN-γ pulses of different concentration. Left: Schematic diagram of the IFN-γ treatment protocol; reporter iBMDMs stimulated with two 1h pulses of IFNγ at 6h interval. Dose of the first pulse varied (100, 10, 5 and 1ng/ml) while the dose of the second pulse was kept constant at 100 ng/ml. Right: Temporal STAT1-tagRFP trajectories in reporter iBMDMs in response to different IFN-γ treatment protocols (as indicated on the graph). Shown are individual nuclear STAT1-tagRFP trajectories as well as ensemble average (in black) for 192, 210, 196 and 204 cells for 100, 10, 5 and 1 ng/ml first dose, respectively, based on duplicated experiments. STAT1-tagRFP fluorescence shown in arbitrary units (a.u.), time in minutes (mins). B. Characteristics of single cell nuclear STAT1 trajectories presented in A. From the left: distributions of peak nuclear STAT1-tagRFP amplitude in response to first (P1) and second pulse (P2). Individual cell data depicted with circles (with mean ± SD per condition) and colour-coded according to treatment protocol. Kruskal-Wallis one-way ANOVA with Dunn’s multiple comparisons test was used to assess differences between groups (*p < 0.05, **p < 0.01, ***p < 0.001, ****p < 0.0001, ns – not significant). Right: Correlation between peak nuclear amplitudes with corresponding Spearman’s correlation coefficients, colour coded according to treatment protocol. C. Distributions of the overall STAT1-tagRFP AUC (over 14h) across treatment protocols. Individual cell data depicted with circles (with mean ± SD per condition) and colour-coded according to treatment protocol. Kruskal-Wallis one-way ANOVA with Dunn’s multiple comparisons test was used to assess differences between groups ****p < 0.0001, ns – not significant).

We hypothesised that the desensitisation is associated with availability of signalling complexes, for example through a depletion of IFNγ R receptors and/or JAK signalling complexes from a total pool available for activation (Lamaze and Blouin 2013; Krause et al. 2006). In this case, in response to sub-saturating doses (which would engage fewer signalling molecules) cells are expected to maintain their responsiveness. In contrast, our data show that even a low dose 1 ng/ml IFN-γ resulted in reduced responses to a subsequent saturated treatment (Fig. 2A). To test this further, reporter iBMDMs were treated with two pulses of IFN-γ at 6 h interval, such that the concentration of the first pulse was 1 ng/ml, while the concentration of the second pulse was varied (100, 10, 5 and 1 ng/ml) (Fig. 3A). We found detectable STAT1 responses to the first and second pulses across all IFN-γ concentrations (Fig. 3B). In response to 1 ng/ml, the peak nuclear STAT1-tagRFP amplitude in the second pulse was significantly lower to that of the first pulse, consistent with pathway desensitisation (Fig. 3B). Similarly, the responses to 5 ng/ml and 10 ng/ml pulses were significantly lower than responses induced by the same dose applied in the first pulse (Fig. 3C). Therefore, these data suggest a model where the first pulse induces a dose-dependent ‘signal threshold’, which subsequently reduces STAT1 responsiveness upon re-exposure to stimulus.

**Figure 3.**
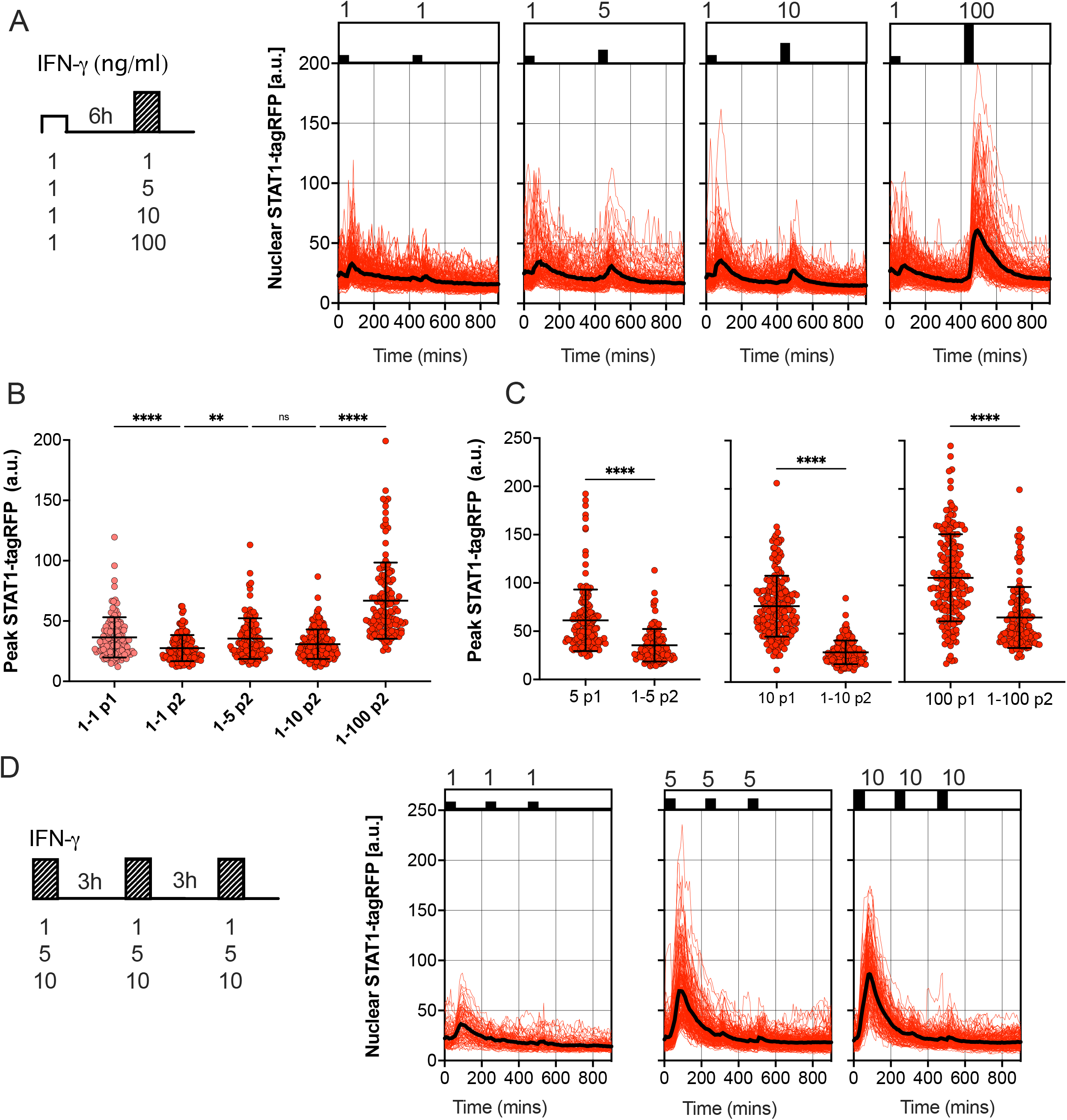
Sub-saturating and high frequency IFN-γ pulses increase signal threshold. A. Single cell analyses of STAT1 responses to IFN-γ pulses of different concentration. Left: Schematic diagram of the IFN-γ treatment protocol; reporter iBMDMs stimulated with two 1h pulses of IFN-γ at 6h interval. Dose of the first pulse kept at 1ng/ml, dose of the second pulse varied (100, 10, 5 and 1ng/ml). Right: Temporal STAT1-tagRFP trajectories in reporter iBMDMs in response to different IFN-γ treatment protocols (as indicated on the graph). Shown are individual nuclear STAT1-tagRFP trajectories as well as ensemble average (in black) for 148, 116, 160 and 132 cells for 1, 5, 10 and 100 ng/ml second dose, respectively, based on duplicated experiments. STAT1-tagRFP fluorescence shown in arbitrary units (a.u.), time in minutes (mins). B. Characteristics of single cell nuclear STAT1 trajectories presented in A. Show is peak nuclear STAT1-tagRFP amplitude in response to first (P1) and second pulse (P2). Individual cell data depicted with circles (with mean ± SD per condition) and colour-coded according to treatment protocol. Kruskal-Wallis one-way ANOVA with Dunn’s multiple comparisons test was used to assess differences between groups (*p < 0.05, **p < 0.01, ***p < 0.001, ****p < 0.0001, ns – not significant). C. Low dose stimulation induces STAT1 desensitisation. Shown are the peak nuclear STAT1-tagRFP amplitudes in response to range of IFN-γ doses (5,10 and 100 ng/ml) in the first (P1) pulse (data from Fig. 2B) compared against the same dose in response to second pulse (P2), when stimulated with 1 ng/ml in the first pulse (data from B). Individual cell data depicted with circles (with mean ± SD per condition) and colour-coded according to treatment protocol. Mann-Whitney pairwise comparisons test was used to assess differences between groups (*p < 0.05, **p < 0.01, ***p < 0.001, ****p < 0.0001, ns – not significant). D. Single cell analyses of STAT1 responses to IFN-γ pulses at 3 h interval. Left: Schematic diagram of the IFN-γ treatment protocol; reporter iBMDMs stimulated with two 1 h pulses of IFNγ at 3 h interval. First and second pulse dose matched, but varied across treatments (1, 5 and 10 ng/ml). Right: Temporal STAT1-tagRFP trajectories in reporter iBMDMs in response to different IFN-γ treatment protocols (as indicated on the graph). Shown are individual nuclear STAT1-tagRFP trajectories as well as ensemble average (in black) for 74, 163 and 138 cells for 1, 5 and 10 ng/ml treatment, respectively, based on duplicated experiments. STAT1-tagRFP fluorescence shown in arbitrary units (a.u.), time in minutes (mins).

Finally, we examined whether the level of desensitisation was related to the timing of IFN-γ stimulation. We subjected reporter iBMDMs to three pulses of IFN-γ at 3 h interval, using matched but different sub-saturating doses (10, 5 and 1 ng/ml) per condition (Fig. 3D). We found that there was no or very little activation in response to the second and third pulse at 3 h interval, regardless of the dose (Fig. 3D). This demonstrates that desensitisation is induced rapidly, before the initial STAT1 response subsides (e.g., in the timescale of 3 h) such that the system is refractory to the same dose (upon re-exposure).

Overall, these data demonstrate that STAT1 desensitisation is dose-dependent, where the level of activation to a stimulation depends on the past (first) IFN-γ dose, revealing a ‘signal memory’ withing the JAK-STAT network. This is consistent with a mechanism, where an initial stimulation sets a ‘signal threshold’, which subsequent treatment must overcome, consequently resulting in reduced responses upon re-exposure. Our data suggest that to elicit a similar signalling response, the dose of IFN-γ upon re-exposure must be higher than that of the initial treatment.

### Type II and type I interferons differentially control JAK-STAT pathway desensitisation

Type II (IFN-γ) and type I IFNs cytokine family (including IFN-α and IFN-β) play distinct functions in the immune response (Barrat, Crow, and Ivashkiv 2019). Both interferons use unique signalling components of the Janus Kinase (JAK)-STAT signalling pathway with a notable exception of signalling adapters, where IFN-γ signals via JAK1 and JAK2 adaptor proteins, while IFN-α and IFN-β engage IFNAR complex and JAK1 and Tyrosine Kinase 2 (TYK2) (Platanias 2005). To provide more insights into control of the JAK-STAT pathway desensitisation, we assayed responses to pulsatile type I and II interferon stimulation. First, we subjected reporter iBMDMs to pulsatile treatment of the combined 50 ng/ml of IFN-α and 50 ng/ml of IFN-β1 (referred hereafter to 100 ng/ml IFN-α/β1). We found that a 1 h pulse induces a single nuclear translocation of STAT1-tagRFP, and the system was refractory to a subsequent pulse of IFN-α/β1 at 6 h interval (Fig. 4AB). This confirms that type I interferons induce complete desensitization of STAT1 signalling in macrophages (Sakamoto et al. 2004). We then treated cells with 1 h pulse of IFN-α/β1 and after 6 h applied a 1 h pulse of IFN-γ (Fig. 4C and D). We found that the population of cells were able to respond to second pulse of IFN-γ, albeit with a reduced amplitude in comparison to a single dose of IFN-γ, demonstrating a partial desensitisation by IFN-α/β1 (Fig. 4E). There was a significant positive correlation (r=0.34) between the 1^st^ and 2^nd^ peak nuclear STAT1 amplitude suggesting that individual cells exhibited similar sensitivity to both stimuli (Fig. 4F). However, when we reversed the order of stimulation cells became refractory to IFN-α/β1, consistent with complete desensitisation by IFN-γ (Fig. 4 C, D and E). These data demonstrate that IFN-γ and IFN-α/β1 differentially regulate JAK-STAT pathway desensitisation, suggesting prioritisation of IFN-γ signalling over IFN-α/β1 through regulatory crosstalk

**Figure 4.**
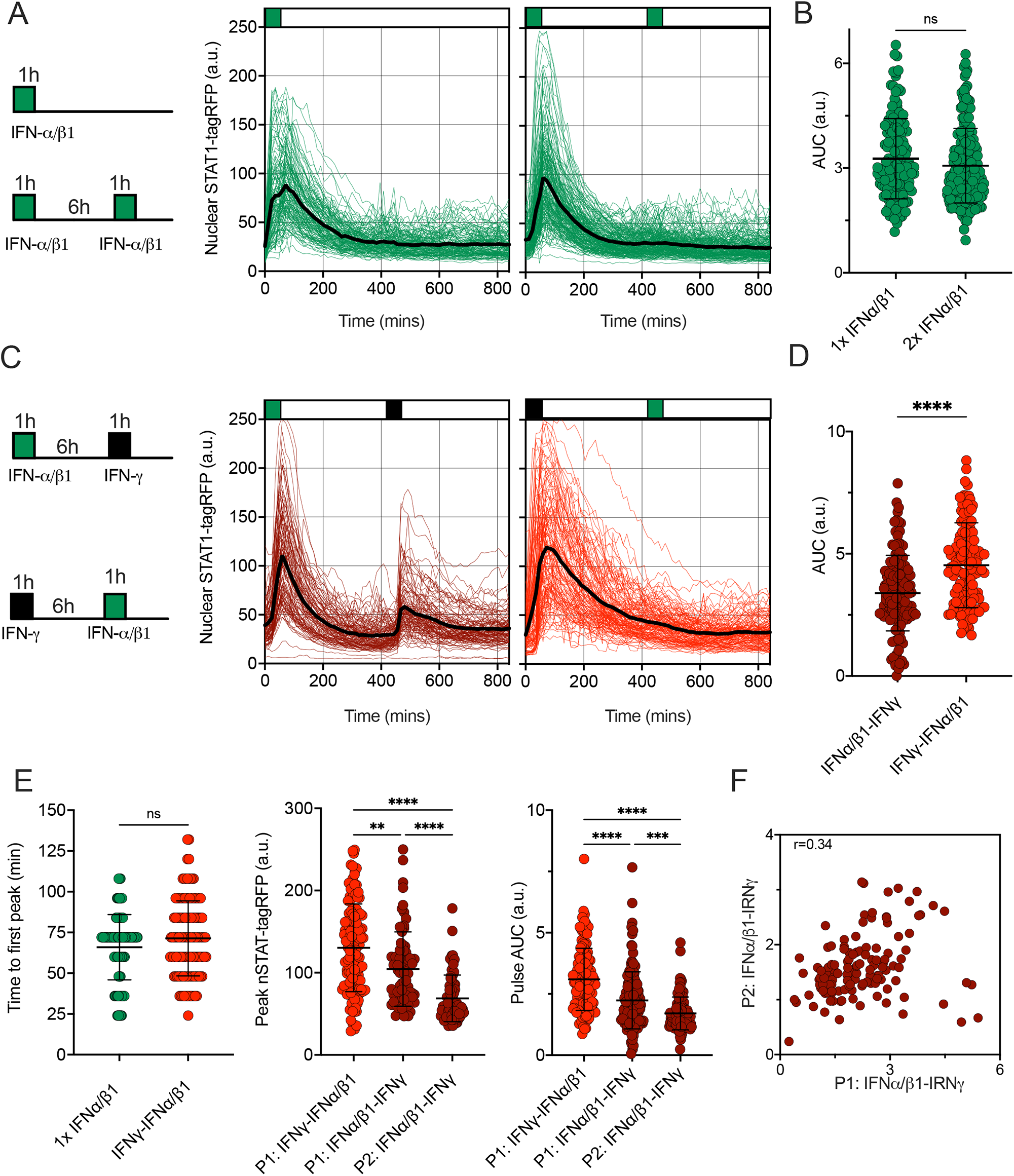
Pathway desensitisation is differentially controlled by type I and II interferons. **A**. Temporal STAT1-tagRFP trajectories in reporter iBMDMs in response to IFN-α/β1 interferon stimulation. Shown are individual nuclear STAT1-tagRFP trajectories as well as ensemble average (in black) for 135 and 193 cells stimulated with 1 h pulse or two 1h pulses at 6h interval of combined 100 ng/ml IFN-α/β1 (50 ng/ml each) from three replicates, respectively. STAT1-tagRFP fluorescence shown in arbitrary units (a.u.), time in minutes (mins). **B**. Distributions of nuclear STAT1-tagRFP AUC in response to IFN-α/β1 pulses (from A). Individual cell data depicted with circles (with mean ± SD per condition). AUC calculated for 14h. Mann-Whitney test was used to assess differences between groups (ns – not significant). **C**. Temporal STAT1-tagRFP trajectories in reporter iBMDMs in response to alternated IFN-α/β1 and IFN-γ stimulation. Shown are individual nuclear STAT1-tagRFP trajectories as well as ensemble average (in black) for 110 and 131 cells stimulated with two alternated 1h pulses at 6h interval of alternated 100 ng/ml IFN-γ and combined IFN-α/β1 (50 ng/ml each) from three replicates, respectively. STAT1-tagRFP fluorescence shown in arbitrary units (a.u.), time in minutes (mins). **D**. Distributions of nuclear STAT1-tagRFP AUC in response to alternated IFN-γ and IFN-α/β1 pulses (from C). Individual cell data depicted with circles (with mean ± SD per condition). AUC calculated for 14h. Mann-Whitney test was used to assess differences between groups (****p < 0.0001). **E**. Characteristics of single cell nuclear STAT1 trajectories presented in A and C. From the left: Distribution of time to first peak, first (P1) and second (P2) peak timing and AUC (Calculated for 7h) under different treatment conditions. Individual cell data depicted with circles (with mean ± SD per condition) and colour-coded according to treatment protocol. Mann-Whitney (for pairwise) and Kruskal-Wallis one-way ANOVA with Dunn’s multiple comparisons test (for three-way comparisons) was used to assess differences between groups (*p < 0.05, **p < 0.01, ***p < 0.001, ****p < 0.0001, ns – not significant). **F**. Correlation between first (P1) and second (P2) nuclear STAT1-tagRFP amplitude in response to alternated IFN-α/β1 and IFN-γ stimulation. Significant Spearman’s correlation coefficient (r) depicted on the graph.

### Desensitisation is regulated post-transcriptionally via STAT1 phosphorylation

Having observed JAK-STAT pathway desensitisation we wanted to understand the underlying molecular mechanisms. IFN-γ exerts its biological function trough binding to the IFNγR1 receptor dimer, which subsequently recruits two chains of IFNγR2 to form the signalling complex (Ivashkiv 2018). IFN-γ uptake initiates internalisation of the IFNγR complex, thus the reduced cell surface expression of the receptor may therefore act as mechanism to limit the level of response (Krause et al. 2006). The quantification of cell-surface receptor expression showed that untreated (control) iBMDMs express a basal level of IFNγR1, which was substantially reduced by continuous 100 ng/ml IFN-γ treatment (Fig. S2). 1 h pulse of 100 ng/ml IFN-γ decreased the expression of IFNγR1 compared to control cells, but most of the receptor was still present on the cell surface. As an additional control, we showed that IFNγR1 expression was not affected by the 100 ng/ml of IFN-α/β1, which specifically bind to its own cognate receptor (Platanias 2005). In terms of IFNγR2 expression, the continuous treatment with IFN-γ substantially reduced IFNγR2 expression at 6h, but neither IFN-γ nor IFN-α/β1 pulses had a substantial impact on the level of expression. Overall, in agreement with live-cell imaging data (Fig. 3A), we conclude that reduced availability of the cell surface IFNγ R expression cannot explain desensitisation of STAT1 signalling.

We next hypothesised that desensitisation is achieved via transcriptional feedback as demonstrated before (Kok et al. 2020; Mudla et al. 2020; Ivashkiv 2018; Yasukawa, Sasaki, and Yoshimura 2000). In particular, IFN-γ -mediated upregulation of SOCS1 is thought to be important for STAT1 responses (Liau et al. 2018). The quantitative smFISH suggested a very low SCOS1 mRNA abundance (up to 10 mRNA molecules per cell) while up-regulation in response to IFN-γ coincided with a change of <1 mRNA molecule on average (Fig. S3A). Therefore, to globally evaluate the role of transcriptional feedback we used RNA polymerase inhibitor Actinomycin D (ActD) (Bensaude 2011) to prevent *de novo* mRNA synthesis and subsequently monitor STAT1 responses via microscopy. In these experiments, reporter iBMDMs were simulated with two 1 h 100 ng/ml IFN-γ pulses at 6 h interval, while treated with ActD either for 2 h before the first pulse or between the two pulses (Fig. 5A). We found that following the ActD treatment STAT1-tagRFP showed a higher level of activity in comparison to control cells, as evident by more prolonged nuclear localisation. Importantly, while ActD treatment did not alter the amplitude of the STAT1 response to the first pulse (Fig. 5B), there were no apparent changes to cell responsiveness as we observed a very limited response to the second pulse. As such, these data suggest that transcriptional feedback may play a role in controlling nuclear localisation of STAT1 (Mowen and David 2000; Begitt et al. 2000), but it is not required to induce desensitisation following IFN-γ stimulation.

**Figure 5.**
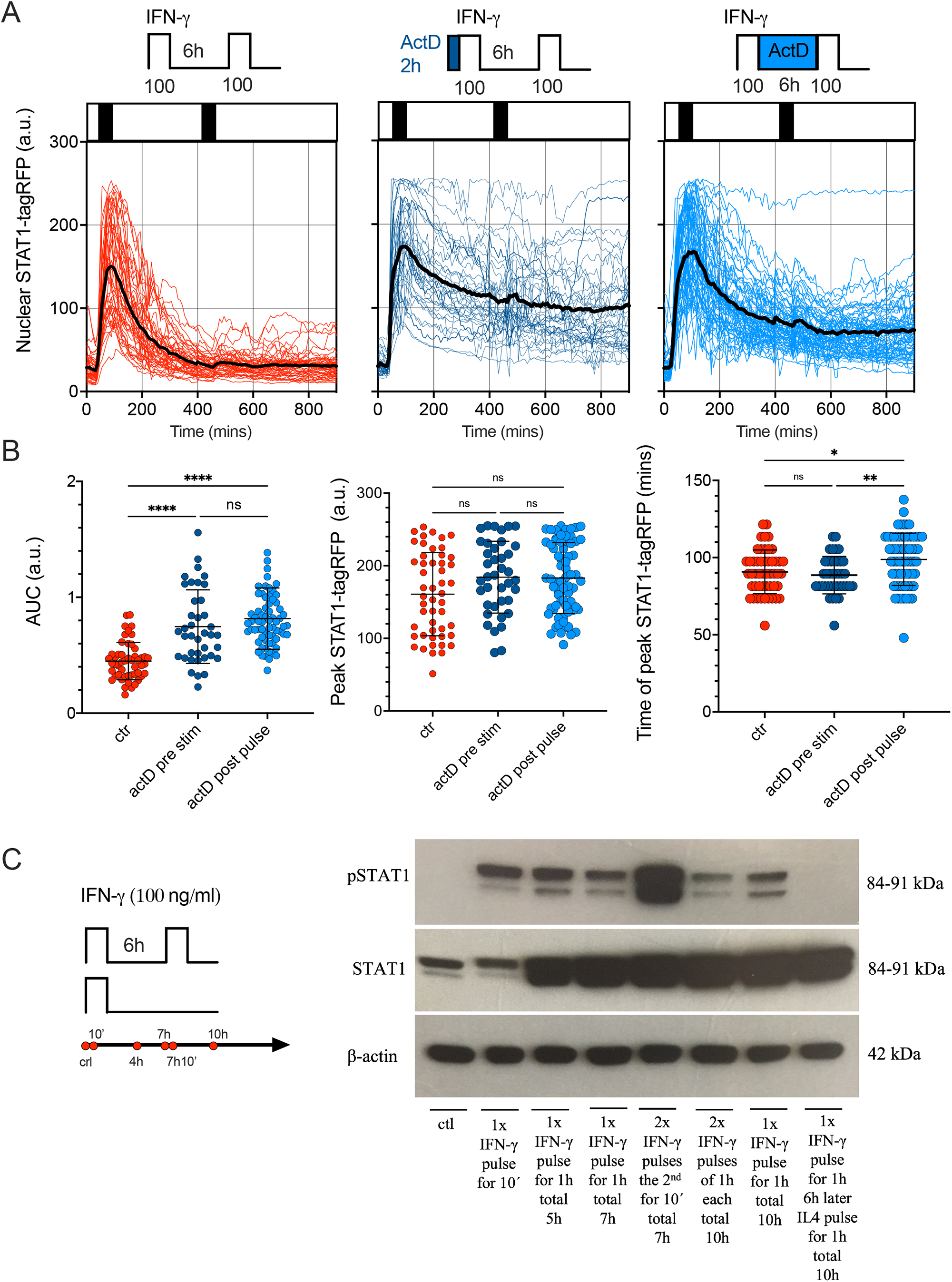
Feedback control of STAT1 desensitisation. A. Single cell analyses of STAT1 responses in presence of transcriptional inhibitors. Reporter cells stimulated with two 1 h pulses of 100 ng/ml IFN-γ (control, left), pre-treated with 5 μg/ml of ActD for 2h before the first pulse (middle) or treated with 5 μg/ml of ActD between the first and second pulse (right). Shown are individual nuclear STAT1-tagRFP trajectories (colour-coded according to treatment protocol) as well as ensemble average (in black) for 50, 39 and 68 cells for control, ActD pre-treatment and ActD treatment between pulses, from two replicates, respectively. STAT1-tagRFP fluorescence shown in arbitrary units (a.u.), time in minutes (mins). B. Characteristics of single cell STAT1 trajectories presented in B. From the left: distributions of the overall AUC, peak amplitude (in response to first pulse), and time to peak (in response to fisrt pulse) under different treatment conditions. Individual cell data depicted with circles (with mean ± SD per condition) and colour-coded according to treatment protocol. Kruskal-Wallis one-way ANOVA with Dunn’s multiple comparisons test was used to assess differences between groups (*p < 0.05, **p < 0.01, ***p < 0.001, ****p < 0.0001, ns – not significant). C. Phosphorylation pattern of STAT1 (Y701) during pulsatile treatment of iBMDMs. Wild type iBMDMs either untreated (ctr) or stimulated with one or two 1h mins pulses of 100 ng/ml IFN-γ at 6h interval. In addition, 100 ng/ml of IL4 was used in the second pulse applied at 6h interval following a pulse of IFN-γ. Samples analysed at 10 mins, 5h, 7h, and 10h after the start of the experiment. B-actin included as a loading control. Schematic diagram represents pulsing protocol and measurement time-points (in red circles). Molecular weight (MW) is shown in kilo Dalton (kDa). Data are representative of two replicates.

Phosphorylation of STAT1 is necessary for cytokine-induced nuclear translocation and regulation of target gene expression (Ivashkiv 2018). Therefore, using Western blotting, we examined phosphorylation patterns of STAT1 tyrosine 701 (Y701), a marker of STAT1 activation in response to pulsatile treatment (Fig. 5C and Fig. S3B). Whereas phosphorylated STAT1 (Y701) was not detected in untreated wild type iBMDMs, *de novo* phosphorylation of STAT1 was highly induced upon 10 mins stimulation with 100 ng/ml of IFN-γ. Phosphorylated STAT1 were also detected in cells treated with a 1 h pulse of 100 ng/ml IFN-γ when examined after 4 h. At 7 and 10 h following the start of the experiment, the STAT1 (Y701) phosphorylation was still maintained, albeit at a lower level (especially at the 10 h time-point). We found an increase of the phosphorylated STAT1 at 10 mins after second IFN-γ pulse (which in part might be due to the upregulated STAT1 protein levels). However, at 4 h after the second pulse (10h from the start of the experiment) the phosphorylation was substantially reduced, and in fact lower than the corresponding response to a single 100 ng/ml IFN-γ pulse (Fig. 5C). As a control we showed that a second pulse of interleukin 4 (IL-4) abolished STAT1 phosphorylation at 10 h. Overall, this data suggested that although STAT1 may be phosphorylated in a response to second IFN-γ pulse, it undergoes a rapid de-phosphorylation, which coincides with lack of nuclear translocation in live-cell microscopy data.

### PTP feedback model of JAK-STAT signalling recapitulates IFN-mediated responses

Our data demonstrates that IFN stimulation results in activation of a post-transcriptional negative feedback, which attenuates STAT1 activation upon re-exposure. This is consistent with action of number of protein tyrosine phosphatases, which are known to inhibit STAT activation (Böhmer and Friedrich 2014). In particular, Tc-PTP (T-cell protein tyrosine phosphatase encoded by PTPN2 gene) was previously found to control STAT1 desensitisation in response to type I and II interferon stimulation (Sakamoto et al. 2004; Heinonen et al. 2009). To quantitatively understand the control of desensitisation we extended our previous model of JAK-STAT signalling (Smieja et al. 2008) and incorporated a new negative feedback due to a putative tyrosine phosphatase PTP (see Fig. 6A for a schematic representation of the mathematical model). The second, positive feedback involved regulation of STAT1 expression, trough STAT1-mediated activation of Interferon Regulatory Factor 1 (IRF1) (Hu et al. 2002; Pertsovskaya et al. 2013). In the model, we assume that IFN-γ and IFN-α/β1 bind their cognate receptors, forming an active receptor complexes, which phosphorylate STAT1 and STAT2 in the cytoplasm (Van Eyndhoven, Singh, and Tel 2021). Phosphorylated STATs (pSTAT1 and pSTAT2) undergo homo- and hetero-dimerization. While the responses to IFN-γ exclusively involve the former, IFN-α/β1 also activates STAT1-STAT2-IRF9 (ISG3) complex (Smieja et al. 2008). The mathematical model representation consisted of 37 differential equations and 65 parameters to recapitulate the ‘average’ behaviour of the JAK-STAT signalling system with a subset of parameters fitted *de novo* (see Tables S2 to S5 for list of variables, ordinary differential equations, fitted parameters and initial conditions). The model was able to accurately recapitulate single-cell STAT1 translocation data for continuous and pulsatile IFN-γ and IFN-α/β1 treatment (Fig. 6 B to D) as well as kinetics of STAT1 mRNA and protein production (Fig. S4A to C).

**Figure 6.**
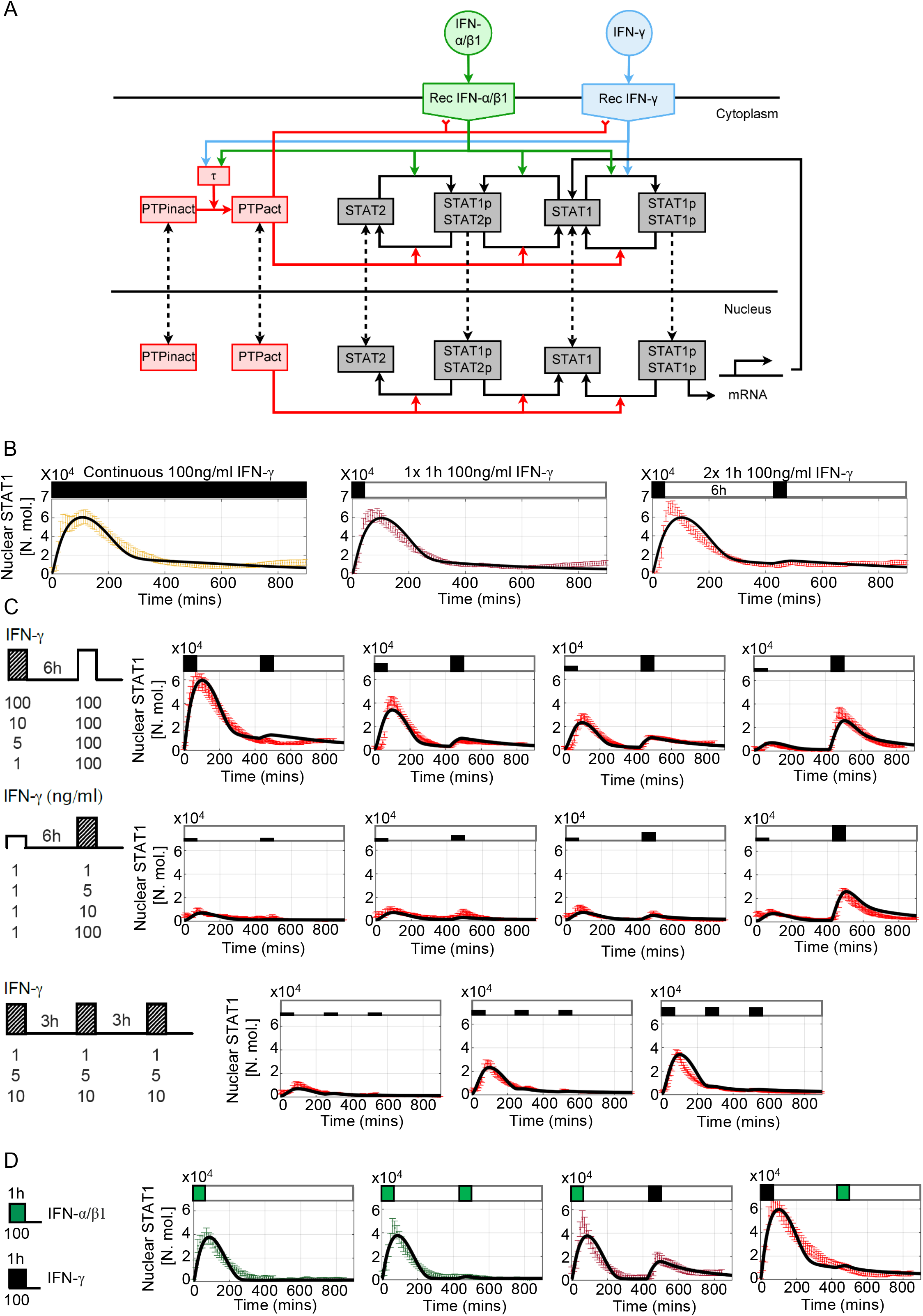
Mathematical model of JAK-STAT pathway desensitisation. **A**. Schematic representation of the JAK-STAT signalling network. IFN stimulation results in receptor binding, activation, and internalisation, leading to STAT1 phosphorylation and translocation to the nucleus. Stimuli-induced putative tyrosine kinase PTP inhibits receptor activity and STAT1 phosphorylation. **B**. Mathematical model recapitulates IFN-γ -induced desensitisation. Shown is the simulated nuclear STAT1 in number of molecules (in black) and scaled experimental data (from Fig. 1B, in colour, shown as mean with 99% confidence intervals). Cells treated with 100 ng/ml of IFN-γ either as continuous (left), one 1h pulse (middle) and two 1 h pulses at 6 h interval. **C**. Model recapitulates responses to different doses and timing of IFN-γ stimulation. Shown are simulated nuclear STAT1 expressed in number of molecules (in black) and scaled experimental data (mean with 99% confidence intervals in red) across different stimulation protocols (as highlighted with schematic diagrams). Top: Two 1 h pulses of IFN-γ at 6h interval as in Fig. 2A, Middle: low dose priming as in Fig. 3A. Bottom: 1 h pulses of IFN-γ at 3 h interval as in Fig. 3C. **D**. Model recapitulates IFN-γ and IFN-α/β1 crosstalk. Simulated nuclear STAT1 expressed in number of molecules (in black) and scaled experimental data (mean with 99% confidence intervals). Cells either treated with 1h pulse of IFN-α/β1, two pulses of IFN-α/β at 6 h interval, or combination of 1 h pulses of IFN-γ and IFN-α/β1 (as in Fig. 4A)

The main regulatory feedback in the model loop involves activation of PTP via the active receptor (Singh et al. 2022). Following previous work, we assumed that activated PTP both inhibits the receptor complex (Simoncic et al. 2002) as well as directly de-phosphorylate STATs (ten Hoeve et al. 2002) leading to dissociation of STAT complexes and nuclear export (Böhmer and Friedrich 2014). *In silico* PTP knockout resulted in complete sensitisation to type I and type II interferons, such that cells exhibited full STAT1 activation in response both IFN-γ or INFα/β1 pulses at 6 h interval (Fig. S4D). While desensitisation relied on both feedback targets, the PTP-mediated de-phosphorylation of STAT1 affected responses more than the inhibition of the receptor complex. In particular, PTP-mediated de-phosphorylation controlled STAT1 nuclear localisation (and recovery to the steady-state) in response to IFN-γ, but not IFN-α/β1 (Fig. S4D). This reflected the differences in the internalised receptor half-life, which has been previously shown to be longer for IFN-γ (Londino et al. 2017) than for IFN-α/β1 (Marijanovic et al. 2006; Kumar et al. 2003) (110 vs 60 mins). Finally, analysis of the model demonstrates that desensitisation was not affected by the level of STAT1 expression, such that a substantial build-up of STAT1 protein had a minor effect on responses to IFN-γ stimulation at 24 h pulsing interval (Fig. S4C).

### Pathway desensitisation renders cells with signal memory of interferon stimulation

Having fitted the new JAK-STAT model to the microscopy data we wanted to systematically investigate mechanisms controlling pathway desensitisation. We assume that in resting cells PTP exists in the inactive form, which upon IFN stimulation undergoes activation via the receptor complex. In response to 100 ng/ml IFN-γ pulse stimulation, PTP activity exhibited saturated non-linear kinetics, characterised by rapid increase to its maximal level at around 6 h (Fig. 7A). At lower IFN-γ concentrations, PTP activation was delayed and reduced over the 800 mins time course, however even the lowest 1 ng/ml dose was able to induce considerable PTP activity. Such a model fit was a consequence of imaging data which demonstrate that 1 ng/ml pulse caused ∼50% reduction in STAT1 response amplitude upon exposure to saturated IFN-γ concentration (Fig. 6C).

**Figure 7.**
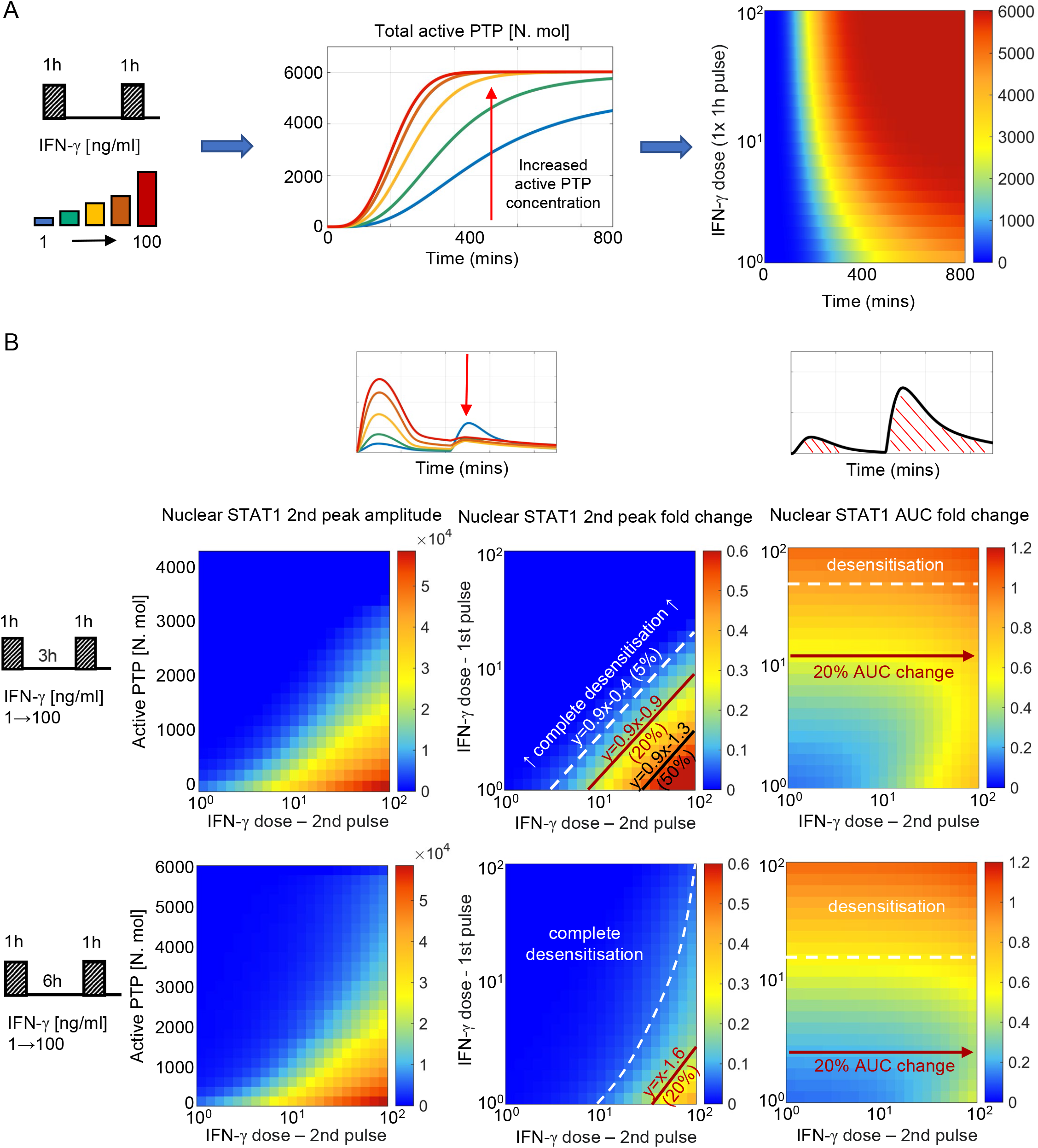
PTP feedback renders cells with signal memory of IFN stimulation. **A**. Dose dependent kinetics of PTP activation: simulations of 1 h IFN-γ pulse across a range of doses (1-100 ng/ml). Middle: Simulated PTP activity over time (in mins) across selected IFN-γ doses (as highlighted on the schematic diagram). Right: heat map of PTP activity (in number of molecules) over range of doses and time (as indicated on the graph). **B**. Signal memory of IFN-γ stimulation: simulations of two IFN-γ pulses applied at 3 (top) and 6 h (bottom) intervals across a range of doses (1-100 ng/ml). Left: Heat-maps of peak nuclear STAT1 upon re-exposure as a function of active PTP level (in number of molecules) and the IFN-γ dose (in log scale) at the time of the 2^nd^ pulse. Middle: Heat maps of peak nuclear STAT1 fold change in response to the 2^nd^ pulse across a range of IFN-γ doses (in log scale). Fold change calculated with respect to the peak nuclear STAT1 in response to 1 h pulse of 100 ng/ml of IFN-γ. Shown in black are relationships corresponding to 5%, 20% and 50% of the response level. Equations depict linear relationships, x and y are concentrations of 1^st^ and 2^nd^ dose, respectively (in log scale). Right: Heat maps of nuclear STAT1 AUC fold change in response to IFN-γ pulses across a range of doses (in log scale). Fold change calculated with respect to the nuclear STAT1 AUC in response to 1 h pulse of 100 ng/ml IFN-γ. In white and red lines shown are relationships corresponding to 10% (desensitisation) and 20% of the AUC with respect to 2^nd^ pulse dose, respectively.

To better understand the relationship between dose of the stimulus, PTP activity and STAT1 translocation kinetics we simulated responses to two IFN-γ pulses for ∼5000 different combinations of doses across 100 ng/ml range (with 1 ng/ml step, Fig. 7B). We find that the different (dose-dependent) levels of PTP activity quantitatively determine whether the system responds to stimulus (at the time of the second pulse) and if it does so, the amplitude of the response. In general, to elicit any signalling response upon re-exposure (defined as the >5% of the amplitude of the single 100 ng/ml IFN-γ pulse) the dose of the 2nd pulse must be substantially higher that the dose of the 1^st^ pulse (Fig. 7B, middle). At 3 h pulsing interval, to achieve 5% response the dose of the second pulse must be ∼3-times larger than the dose of the 1^st^ pulse, while ∼8- and ∼30-times larger concentrations are required to achieve 20 % and 50% response, respectively. As the level of PTP activity increases with time, the sensitivity to second pulse is reduced, such that at 6 h, the dose of the second pulse must be at least ∼35 times higher than the dose of the first pulse to elicit a 20% response, while 50% response can only be achieved following a 1 ng/ml stimulation. Notably, the observed relationships between the doses are linear (apart for the 5% response for the 6 h pulsing interval), while the same level of response to the 2^nd^ pulse can be achieved by combinations of IFN-γ concentration, suggesting that the system may effectively respond to the relative fold changes of input concentration. Subsequently, we investigated the overall sensitivity of STAT1 response to IFN-γ pulses (Fig. 7B, right). We found that the AUC of the nuclear STAT1 in response to two pulses (at 3 and 6 h pulsing intervals) was inherently restricted; that is the overall responses was not higher than the AUC of the single 1 h 100 ng/ml pulse (calculated over the same time interval). For concentrations above >16 ng/ml of IFN-γ at 6 h pulsing intervals (and >50 ng/ml for 3 h) any further stimulation had a minor effect on the overall response (i.e., the relative AUC changes <10% with respect to the 2^nd^ pulse dose), but the system exhibited sensitivity the 1^st^ pulse dose. In turn, at lower concentration (<2.5 ng/ml for 6 h and <12 ng/ml for 3 h intervals) the overall AUC exhibited increased sensitivity to the 2^nd^ pulse (i.e., >20% AUC change), in particular high IFN-γ concentrations. This demonstrates that while maintaining the overall dose-dependency, the PTP feedback restricts the temporal JAK-STAT signalling response to that of a single transient and saturated IFN-γ input.

Finally, we investigated the role of the PTP feedback in the crosstalk between IFN-γ and IFN-α/β1 pulses. In agreement with experimental data, the mathematical model demonstrated that IFN-α/β1 stimulation induced a lower STAT1 translocation (Fig. 4E), which resulted in a lower PTP activity, in comparison to matching doses of IFN-γ (Fig. S5A). Consequently, PTP activity induced via IFN-α/β1 was not sufficient to inhibit responses upon to IFN-γ stimulation. We found that ∼1.5-fold change in IFN-γ dose (comparing to that of IFN-α/β1) was required to elicit a 5% response and ∼4-fold change for a 20% of the response upon re-exposure (Fig. S5B). Consequently, the overall AUC of nuclear STAT1 exhibited more sensitivity to IFN-γ (defined as 20% AUC change) than to IFN-α/β1, in cells stimulated with IFNα/β1 in the first pulse. In turn, the level IFN-γ -induced STAT1 activation was higher, and consequently higher PTP levels almost completely inhibited responses to subsequent IFN-α/β1 stimulation, resulting in complete desensitisation. Overall, these analyses demonstrate that the PTP feedback renders cells with signal memory by responding to the relative fold changes of the IFN concentration and discriminate different temporal patterns of type I and type II interferon stimulation.

## Discussion

Here we provide a new quantitative understanding of JAK-STAT signalling in response to temporal type I and type II IFNs in innate immune macrophages. Using live-cell microscopy to follow intracellular dynamics of STAT1 localisation we demonstrate that responses to interferon stimulation are tightly controlled trough the desensitisation of the JAK-STAT pathway. We demonstrate that a brief 1 h stimulation with saturating concentration of IFN-γ elicit quantitatively similar response to that of continuous stimulation, resulting in a complete inhibition of the signalling and gene expression responses for up to 24 h. JAK-STAT signalling is regulated at multiple levels, including the receptor availability (Bach et al. 1995) as well as regulatory feedback (Kok et al. 2020; Mudla et al. 2020; Ivashkiv 2018; Yasukawa, Sasaki, and Yoshimura 2000; Liau et al. 2018; Smieja et al. 2008). For example, desensitisation to IFN-α stimulation is controlled trough the transcriptional feedback due SOCS1 and USP18 (Kok et al. 2020; Mudla et al. 2020). Here we demonstrate that in macrophages desensitisation involves attenuation of STAT1 phosphorylation resulting in lack of a nuclear translocation. Importantly *the novo* mRNA synthesis (and thus transcriptional feedback) and IFN-γ R availability cannot explain the IFN-γ -mediated responses. Our data is consistent with a post-transcriptional feedback mechanisms involving the activation of the protein tyrosine phosphatase activity. The current knowledge demonstrates that multiple PTPs may regulate JAK-STAT signalling, including transmembrane CD45 and PTPε as well as intracellular Tc-PTP, PTP1B and SHP-1, which attenuate STAT phosphorylation at different levels (Böhmer and Friedrich 2014). Based on this knowledge we extended our previous mathematical model of IFN-β-mediated JAK-STAT signalling system (Smieja et al. 2008) to demonstrate that PTP feedback recapitulates detailed imaging data on the dose, timing and type of interferon stimulation. Our mathematical model likely captures the combined effect of multiple PTPs involved, however it suggests that neither the ability of PTP to block receptor complex nor dephosphorylate STAT1 is sufficient alone to recapitulate observed responses. We believe, that considering the current knowledge of IFN signalling, the Tc-PTP could be a primary feedback mechanism, as it was previously shown to regulate sensitivity to IFN-β and IFN-γ stimulation (Sakamoto et al. 2004). Primary human fibroblast exhibited partial desensitisation to IFN-β after 16 h IFN-β treatment as well as to subsequent IFN-γ stimulation (at doses <5 ng/ml applied 6 h after initial stimulation) and the attenuation of STAT1 phosphorylation was present in wild type, but not Tc-PTP knockout cells (Sakamoto et al. 2004). However, surprisingly little is known about regulation of Tc-PTP by IFNs, but recent work demonstrates a direct molecular interaction between a cognate receptor (integrin 1α in the context of cell adhesion) activates Tc-PTP by disrupting the autoinhibitory C-terminal tail of the kinase (Singh et al. 2022). Further work is required to understand the kinetics of Tc-PTP activity, which we predict lasts beyond the internalised receptor half-life (>24h) as well as to understand potential contributions of other PTPs and in fact other STAT molecules in this process. In the broader context, the quantitative understanding of PTP regulation might provide important insights into control of IFN signalling during immune responses as well as oncogenic signalling (Pike and Tremblay 2016).

Desensitisation is a key mechanism that prevents prolonged out-of-control activity to chronic stimulation and/or limit responses upon re-exposure to the same stimulus, for example in the toll-like receptor system (Morris, Gilliam, and Li 2014; Buckley, Wang, and Redmond 2006).

Our quantitative imaging data and mathematical modelling demonstrate that through pathway desensitisation, the overall JAK-STAT response is restricted, such in response to repeated cues cells cannot produce more activity than to a single 1 h pulse with a saturated IFN-γ concentration. Stimulation with sub-saturating IFN-γ concentrations resulted in partial STAT1 desensitisation, through a dose- and stimulus-dependent negative feedback, and consequently the system’s responses upon re-exposure depended on the dose of the initial stimulation and its timing. We found that to elicit a signalling response upon re-exposure, the concentration of IFN-γ at 3 and 6 h must be several fold larger than that of the initial stimulation. This effectively means that the JAK-STAT system becomes refractory following stimulation with medium and high IFN-γ doses but retains reduced sensitivity to lower concentrations (<10 ng/ml). These analyses demonstrate that through post-transcriptional PTP feedback, desensitisation of JAK-STAT signalling renders cells with signal memory by responding to relative fold changes in IFN concentration. In comparison, the NF-κB system can detect absolute increases in cytokine concentration during relatively short (<2h) time intervals (Son et al. 2021). Other systems use receptor availability to detect temporal changes in stimulus, for example, relative (fold) changes in early growth response (EGR) protein concentration (Lyashenko et al. 2020). Interestingly, we demonstrate that PTP feedback may also distinguish different patterns of type I and II interferon stimulation, such that cells refractory to IFN-α/β1 are sensitive to IFN-γ, but not *vice versa*. The type I and II IFNs use unique receptor complexes, thus the overlapping receptor-associated adapters and target STAT activation may result in a functional crosstalk (Takaoka et al. 2000). Here we demonstrate that IFN-γ and IFN-α/β1 elicit not only different STAT1 translocation profiles, but also different levels of desensitisation, where cells exhibit more sensitivity to IFN-γ than to IFN-α/β1 (per dose), leading to increased PTP activity in response to IFN-γ and subsequently suppression of IFN-α/β1 responses upon re-stimulation. Currently, our model provides a limited insight into differences between IFN-γ and IFN-α/β1 signal transduction as we assumed the same rate kinetics except of receptor half-lives (Londino et al. 2017; Marijanovic et al. 2006; Kumar et al. 2003). However, it will be important to understand signal specific mechanisms and in particular receptor specificity for different PTPs (Böhmer and Friedrich 2014). We suggest that prioritisation of IFN-γ signalling might reflect different roles during pathogen infection, tissue specificity and timing (Ivashkiv 2018) and reflect specific interactions with pathogens, for example *L. monocytogenes*, where type I and type II interferons induce opposite effects in terms of host susceptibility (Rayamajhi et al. 2010).

Previous work demonstrates the involvement of positive feedback in JAK-STAT signalling; low-dose IFN-γ sensitised human PMBCs to IFN-α stimulation trough upregulation of JAK-STAT signalling molecules including STATs (Hu et al. 2002; Pertsovskaya et al. 2013; Lehtonen, Matikainen, and Julkunen 1997). JAK-STAT pathway desensitisation was also shown to depend on the dose of the treatment in hepatocytes, where a 24 h pre-treatment with ∼50 pg/ml of IFN-α resulted in increased responsiveness upon re-exposure, while 25 ng/ml IFN-α induced desensitisation (Kok et al. 2020). In this work we only use concentrations of 1 ng/ml and above since we found that lower concentrations do not induce robust STAT1 translocations via imaging, However, we expect that the specific effects might reflect differences between (immune vs non-immune) cell types. Our model captured population-level data on the IFN-γ -induced STAT1 up-regulation in our system, which we showed had a limited effect on STAT1 responses in the model at least up to 24 h after stimulation. It is possible that in a longer term (while PTP activity subsides) and the STAT levels increase substantially the JAK-STAT system becomes more sensitive to stimulation. Finally, recent analyses suggest digital activation of STAT1 to IFN-γ, where only a fraction of mouse embryonic fibroblasts responding with STAT1 phosphorylation to a particular dose (Topolewski et al. 2022). Our data, in agreement with previous work on the NF-κB system, suggest that macrophages exhibit analogue encoding (Sung et al. 2014; Bagnall et al. 2018), where the dose controls the amplitude of the response, rather than a fraction of responding cells. However, it would be important to understand the variability of key STAT target genes as it may provide the insight into the overall transcriptional control of IFN responses (Bagnall et al. 2020). Overall, our analyses demonstrate IFN mediated signalling responses to pulsatile cues are tightly constrained, which we believe facilitates the need within the immune system to control pathological interferon signalling.

## Author contributions

EK developed cells for imaging, collected, and analysed data. MK and JS developed mathematical model. JB assisted with cell line development and imaging analyses. DS assisted with imaging analyses. WM, DR, SB and PP provided supervision and conceptualisation. PP with assistance of EK and MK wrote the manuscript. All authors read and approved the final manuscript.

## Acknowledgments

We would like to thank Dr Kevin Couper for reagents.

## Data Availability Statement

Tabularized manuscript data in provided in Table 1, mathematical models generated in this study will be available from Github repository (https://github.com/ppaszek/Jak-STAT-signal-memory) upon publication.

---

**Movie 1. Continuous stimulation with 100 ng/ml IFN-γ**. Confocal microscopy movie of iBMDM cells expressing STAT1-tagRFP (red channel), Venus-STAT6 (yellow channel) and AmCyan-H2B (cyan channel). The time from the start of the experiment is depicted in min. Cells stimulated at time 0 mins.

**Table 1. Tabularised manuscript data**.

