## Supplementary material for "Post-transcriptional regulatory feedback encodes JAK-STAT signal memory of interferon stimulation": Suplementray information

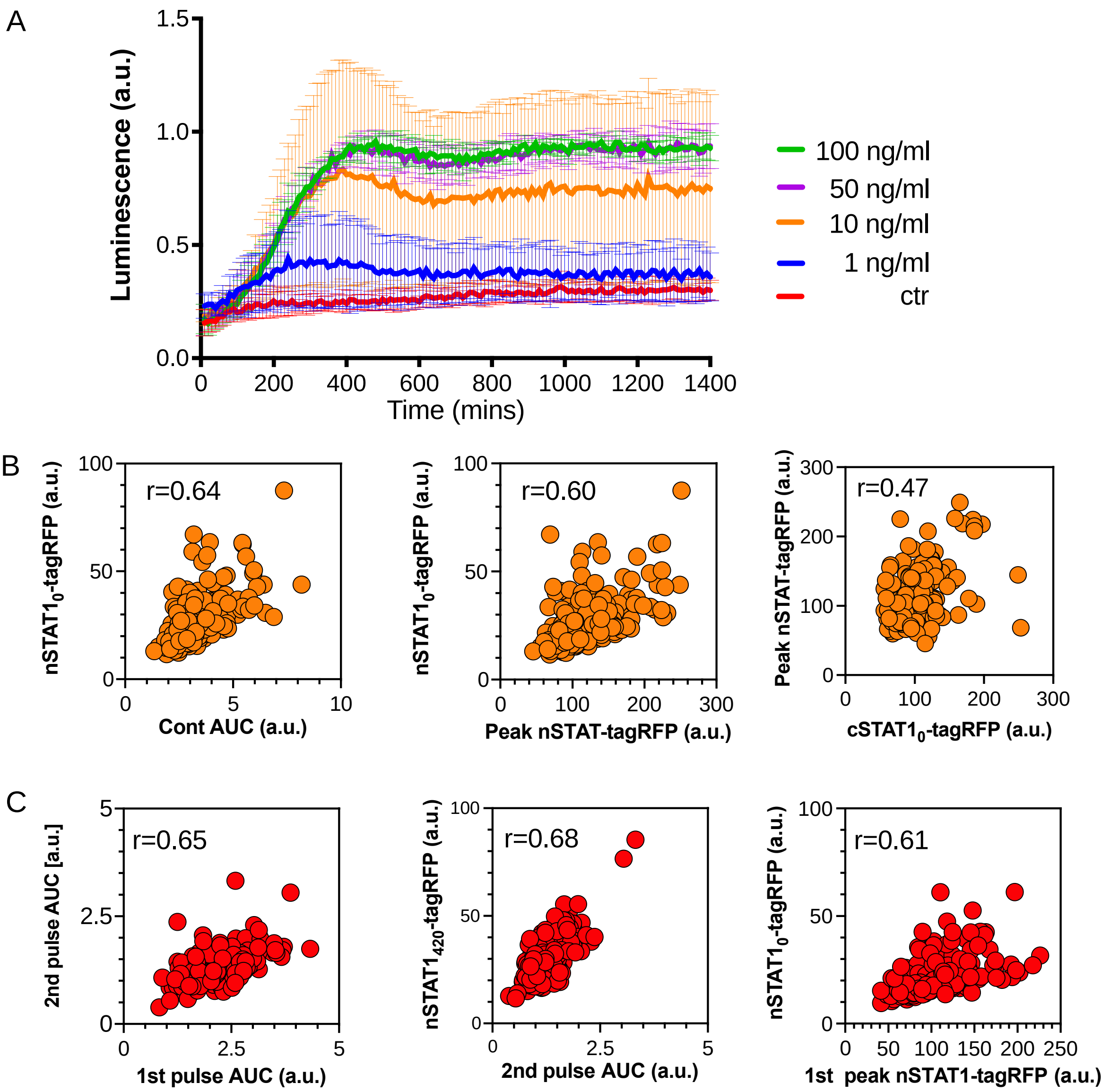


**Figure S1. Characteristics of STAT1 activation. A.** Live-cell luminometry analysis of GAS-mediated expression in response to range of IFN-γ doses. Shown are the average (and SDs) GAS-luc trajectories in iBMDMs, in response to continuous 1, 10, 50 and 100 ng/ml of IFN-γ treatment compared to unstimulated cells (ctr). Luminescence in arbitrary units (a.u.), time in minutes. Data obtained from three replicates. **B.** Analysis of the response to continuous 100 ng/ml IFN-γ from Fig. 1B; AUC vs. nuclear STAT1-tagRFP at t=0 mins and peak nuclear STAT1-tagRFP (nSTAT1-tagRFP) vs. nuclear STAT1-tagRFP at t=0 mins (nSTAT1$_0$-tagRFP); peak nuclear STAT1-tagRFP vs. cytoplasmic STAT1-tagRFP at t=0 mins (cSTAT1$_0$-tagRFP). **C**. Analysis of the response to two 1 h pulses of 100 ng/ml IFN-γ Fig. 1B; AUC in response to first vs. second pulse calculated over 420 mins each, AUC in response to second pulse vs. nuclear STAT1-tagRFP at t=420 mins; peak nuclear STAT1-tagRFP in response to first pulse vs. nuclear STAT1-tagRFP at t=0 mins. Significant (positive) Spearman's correlation coefficient (r) depicted on each panel.

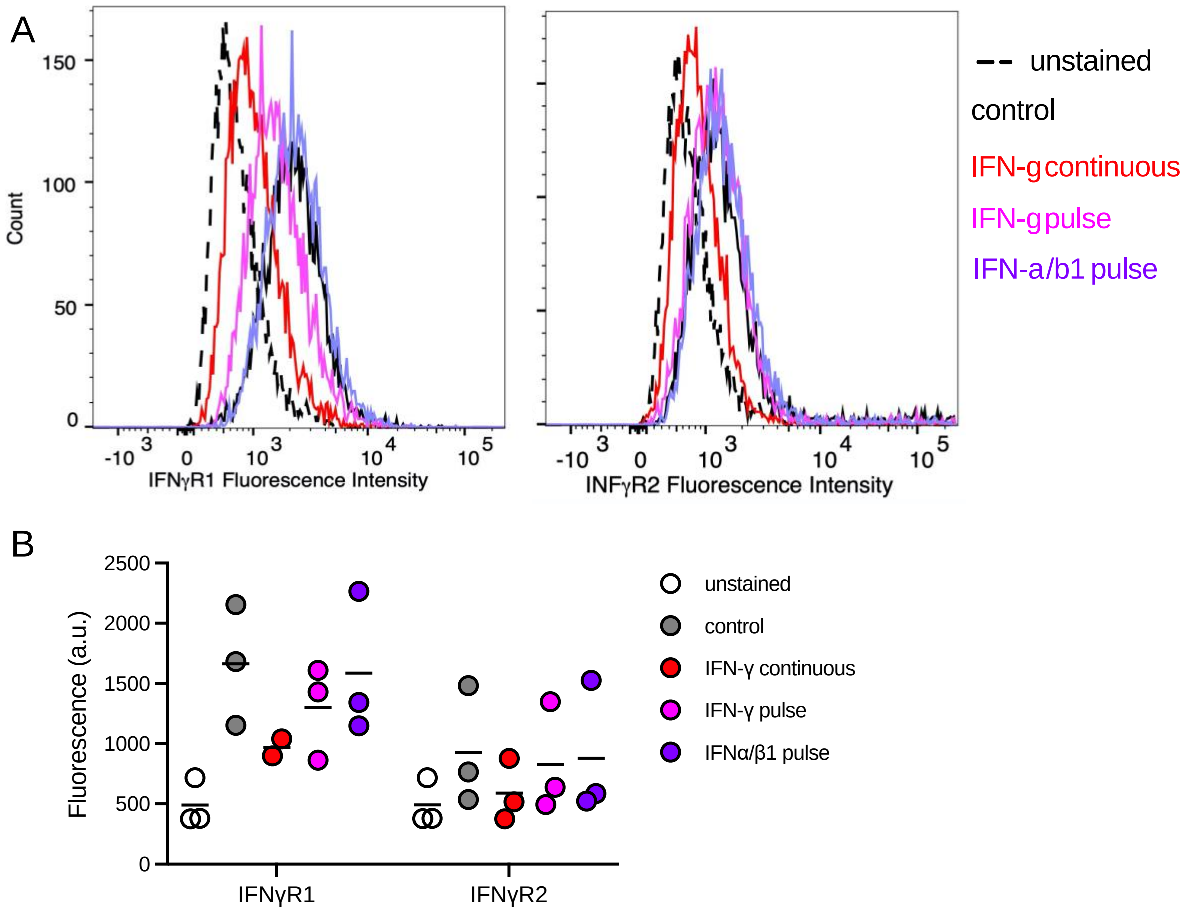


**Figure S2. Expression of IFNγ-R1 and IFNγ-R2 in iBMDMs in response to type I and II IFNs. A**. IFNγ-R1 and IFNγ-R2 cell surface expression in wild-type iBMDMs unstained (dashed line) and untreated (in black solid line) as well as treated with continuous 100 ng/ml IFN-γ (in red), one 1h pulse of 100 ng/ml IFN-γ (pink colour) and a combined 1h pulse of 50 mg/ml of IFN-α and IFN-β1 (in purple) and measured at 7h from the beginning of the experiment. Shown are histograms (number of cells (x1000)) of fluorescent intensity in arbitrary units (x1000). **B**. Geometric mean fluorescence intensity of IFNγ-R1 and IFNγ-R2 staining for data in A. Shown are individual geometric means as well as the mean of three replicates.

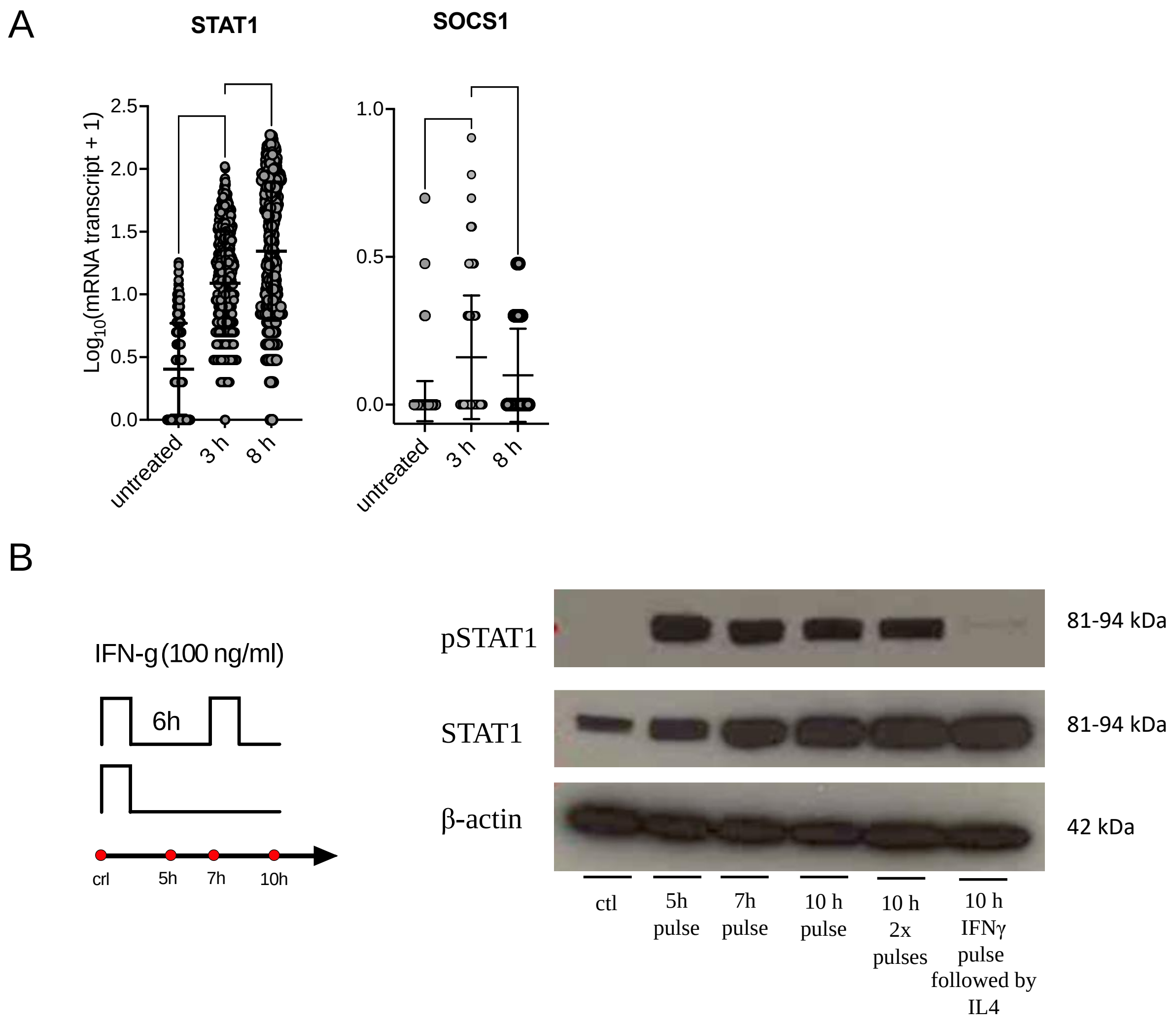


**Figure S3. Regulation of STAT1 and SOCS1 by IFN-γ. A**. smFISH analysis of STAT1 and SOCS1 mRNA expression upon IFN-γ treatment. Wild-type iBMDMs either untreated (ctrl, N=551) or stimulated with IFN-γ for 3 h (N=574) and 8 h (N=567). Number of mRNA transcripts expressed in $\log_{10}$(mRNA transcripts + 1). Circles correspond to individual cell data with population averages (and SDs) in black (based on duplicated experiment). Results of Dunn's multiple comparisons test for statistical difference between the different time points are presented in the top (**** - $p \leq 0.0001$). **B.** Phosphorylation pattern of STAT1 (Y701) during pulsatile treatment of iBMDMs. Wild type iBMDMs either untreated (ctr) or stimulated with one or two 1h mins pulses of 100 ng/ml IFN-γ at 6h interval. In addition, 100 ng/ml of IL4 was used in the second pulse applied at 6h interval following a pulse of IFN-γ. Samples analysed at 5h, 7h, and 10h after the start of the experiment. B-actin included as a loading control. Schematic diagram represents pulsing protocol and measurement time-points (in red circles). Molecular weight (MW) is shown in kilo Dalton (kDa). Data are representative of two replicates.

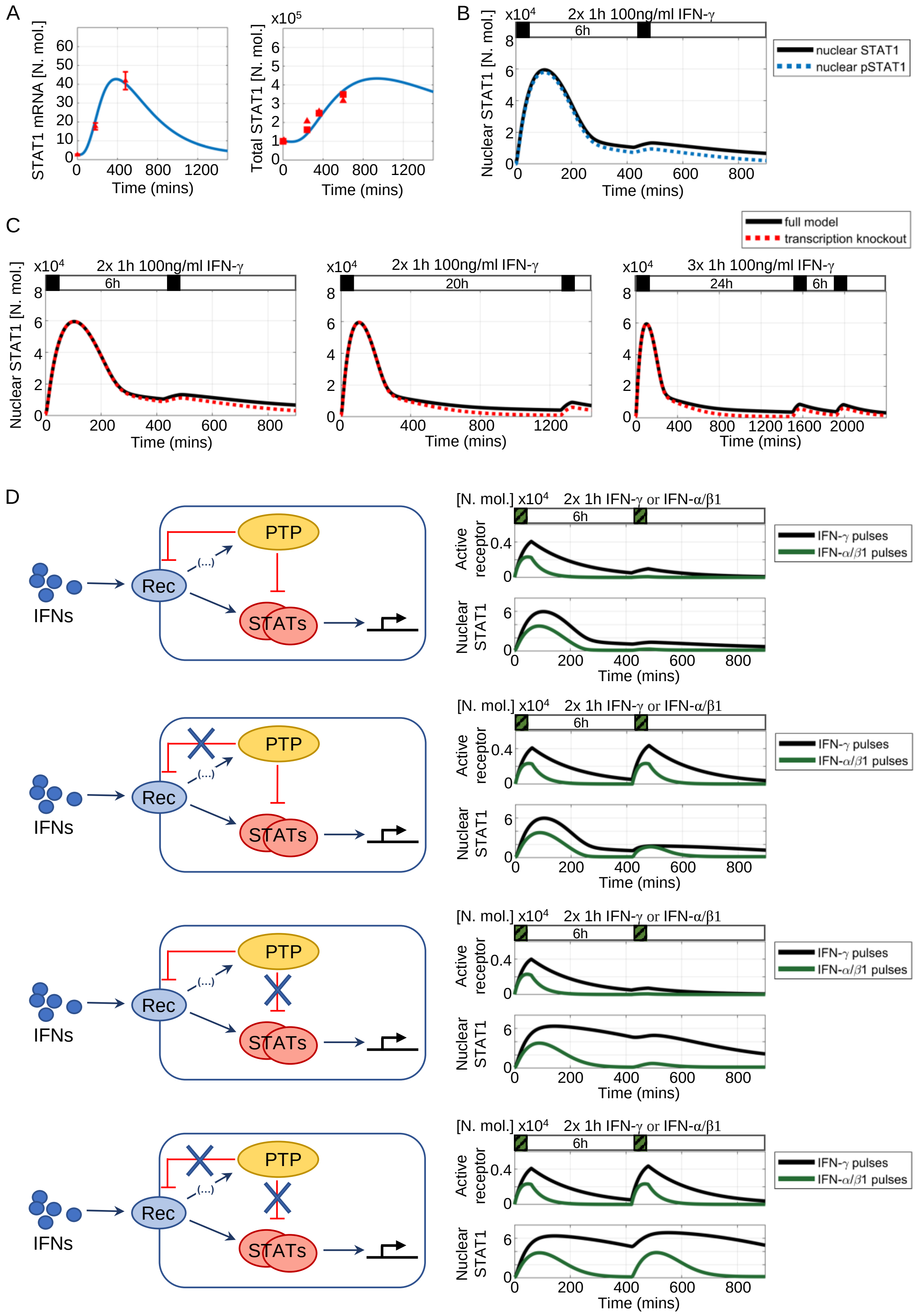

A
STAT1 mRNA [N. mol.]
Time (mins)
x10^5
Total STAT1 [N. mol.]
Time (mins)
B
x10^4
2x 1h 100ng/ml IFN-γ
6h
Nuclear STAT1 [N. mol.]
nuclear STAT1
nuclear pSTAT1
Time (mins)
C
full model
transcription knockout
x10^4
2x 1h 100ng/ml IFN-γ
6h
Nuclear STAT1 [N. mol.]
Time (mins)
x10^4
2x 1h 100ng/ml IFN-γ
20h
Time (mins)
x10^4
3x 1h 100ng/ml IFN-γ
24h
6h
Time (mins)
D
IFNs
Rec
PTP
(...)
STATs
[N. mol.] x10^4
2x 1h IFN-γ or IFN-α/β1
6h
Active receptor
Nuclear STAT1
Time (mins)
IFN-γ pulses
IFN-α/β1 pulses

**Fig. S4. Feedback regulation in the JAK-STAT network. A.** Temporal regulation of STAT1 mRNA (left) and protein (right) level. Shown in the comparison between model simulations (blue line) and experimental data (in red) in cells responding to continuous 100 ng/ml IFN-γ treatment. mRNA data from Fig. S3B shown as mean (with 99% confidence intervals) across times points. Total cellular protein levels obtained from quantification of immunoblotting data in Fig. 5C (triangles) and Fig. S3C (squares). **B**. Simulated STAT1 trajectories in response to two 1 h pulses of IFN-γ at 6 h interval. Shown are nuclear STAT1 (in black) and pSTAT1 (in blue). **C**. Simulated nuclear STAT1 trajectories in response to pulsatile IFN-γ stimulation. Shown are responses of WT cells (in black) in comparison to inducible STAT1 transcription knock out (in red). STAT1 transcription knockout was performed by setting inducible transcription rate vs1 to 0. Cells simulated with two or three 1 h pulses of 100 ng/ml IFN-γ at different intervals (as indicated on the graph). **D.** Contribution of PTP-mediated receptor complex inhibition and dephosphorylation of pSTAT1 on the nuclear STAT1 in the mathematical model. Cells stimulated with two 1 h pulses of 100 ng/ml IFN-γ (in black) or 100 ng/ml IFN-α/β1 (in green). Shown is the comparison between WT cells and cells with individual (or both) feedback targets removed as indicated with schematic diagrams. Removal of PTP-mediated receptor complex inhibition and dephosphorylation of pSTAT1 modelled by setting kPTPss =0 (PTP|STAT1 complexes association rate) and/or PTPinRa=0 and PTPinRg=0 (PTP-mediated inhibition of active IFN-α/β1 and IFN-γ receptor, respectively) in the mathematical model.

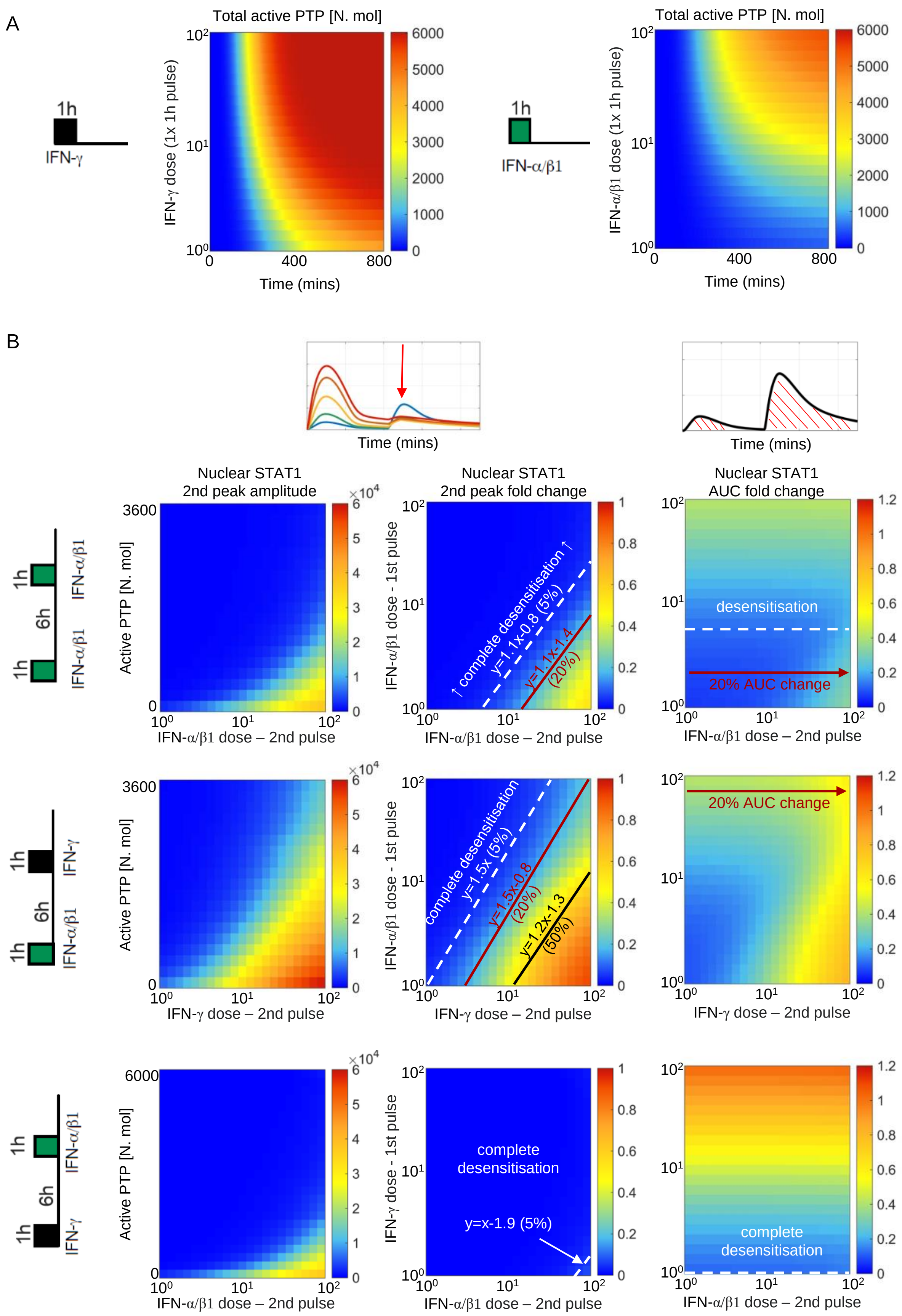
A
1h
IFN-γ
Total active PTP [N. mol]
IFN-γ dose (1x 1h pulse)
Time (mins)
1h
IFN-α/β1
Total active PTP [N. mol]
IFN-α/β1 dose (1x 1h pulse)
Time (mins)
B
Time (mins)
Time (mins)
Nuclear STAT1
2nd peak amplitude
Nuclear STAT1
2nd peak fold change
Nuclear STAT1
AUC fold change
1h
6h
1h
IFN-α/β1
IFN-α/β1
Active PTP [N. mol]
IFN-α/β1 dose – 2nd pulse
IFN-α/β1 dose - 1st pulse
complete desensitisation
y=1.1x-0.8 (5%)
y=1.1x-1.4
(20%)
IFN-α/β1 dose – 2nd pulse
desensitisation
20% AUC change
IFN-α/β1 dose – 2nd pulse
1h
6h
1h
IFN-γ
IFN-α/β1
Active PTP [N. mol]
IFN-γ dose – 2nd pulse
IFN-α/β1 dose - 1st pulse
complete desensitisation
y=1.5x (5%)
y=1.5x-0.8
(20%)
y=1.2x-1.3
(50%)
IFN-γ dose – 2nd pulse
20% AUC change
IFN-γ dose – 2nd pulse
1h
6h
1h
IFN-α/β1
IFN-γ
Active PTP [N. mol]
IFN-α/β1 dose – 2nd pulse
IFN-γ dose - 1st pulse
complete
desensitisation
y=x-1.9 (5%)
IFN-α/β1 dose – 2nd pulse
complete
desensitisation
IFN-α/β1 dose – 2nd pulse

**Fig. S5. Model analysis of the IFN-γ and IFN-α/β1 crosstalk.** A. Dose dependent kinetics of TPT activation: simulations of 1 h IFN-γ or IFN-α/β1 pulse across a range of doses (1-100 ng/ml). Shown are heat maps of PTP activity (in number of molecules) over range of doses and time (as indicated on the graph). **B.** Simulations of 1 h pulse of IFN-α/β1, two pulses of IFN-α/β at 6 h interval, or combination of 1 h pulses of IFN-γ and IFN-α/β1 (as in Fig. 4A) across a range of doses (1-100 ng/ml). Left: Heat-maps of peak nuclear STAT1 upon re-exposure as a function of active PTP level (in number of molecules) and the 1st pulse dose (in log scale) at the time of the 2nd pulse. Middle: Heat maps of peak nuclear STAT1 fold change in response to the 2nd pulse across a range of doses (in log scale). Fold change calculated with respect to the peak nuclear STAT1 in response to 1 h pulse of 100 ng/ml of IFN-γ. Shown are relationships corresponding to 5%, 20% and 50% of the response level. Equations depict linear relationships, x and y are concentrations of 1st and 2nd dose, respectively (in log scale). Right: Heat maps of nuclear STAT1 AUC fold change in response to IFN pulses across a range of doses (in log scale). Fold change calculated with respect to the nuclear STAT1 AUC in response to 1 h pulse of 100 ng/ml IFN-γ. Shown are also relationships corresponding to 10% (desensitisation, in white) and 20% of the AUC (in red) with respect to 2nd pulse dose.

| STAT1 | SOCS1 |
|---|---|
| tgaagctcgaaccactgtga gatttccatggggaaactgt<br>tggtcgcaaacgagacatca tgtgagaggaggtcatggaa<br>gagaaaagcggctgtactgg cgtatgttgtgctgcaacaa<br>tatcctggagattacgcttg atctgtacgggatcttcttg<br>gggcctgattaaatctttgg aaacatcttgtggagcagca<br>ctgagtgagctcgatggaat cacgagctcgtcattaatca<br>tgcaacaatggtgaaccagc ctttttaagctgctgacgga<br>taatagggtcgggctcatag gaaggagctctgaatgagct<br>gtctcagcttgacagtgaac tgcgtacccaagatgttgaa<br>agtgaagttcttcggtgaca cacgacaggaagagaggtgg<br>gttgtaccacaggatagacg ggggttcaggaagaaggaga<br>aactcaacacctctgagagc ctctcttggtgactgatgaa<br>tggaataagaccatcagggc tgtcaatccaaggccagaag<br>gcttcttaatgagctctagg gctctcactgaatctaagca<br>cacccatgtgaatgtgatgg atggaagtcaggttcacctc<br>ccatgactttgtagttgcga attctctggtatgttctcgg<br>tggggtacagatacttcagg aaaggcgtggtctttgtcaa | tgctaccatcctactcgag gagatcgcattgtcggctg<br>ggacgaggagggctctgac cgctggcgaggacgaagac<br>aaggtgcggaagtgagtgt cggtaatcggagtgggagc<br>aatagaagccgcaggcgtc actgtcgcgcaccaagaag<br>gaagaagcagttccgttgg aagccatcttcacgctgag<br>ctggaagtgcacgcggatg agcagctcgaaaaggcagt<br>cgatgcgctggcgacacag taccgggttaagagggatg<br>ggaactcaggtagtcacgg cggtcagatctggaagggg<br>acttaatgctgcggcacag agaagtgggaggcatctca<br>tgagaggtgggatgaggtc acaagctgctacaaccagg<br>gagtggaattcaggtcctg ccctggtttgtgcaaagat<br>aatgaagccagagaccctc ggatattctgcacagcaga<br>attacctaaactggctgta |

**Table S1. Custom smFISH probe sets (5’-3’).**

| Variable: | Description: |
|---|---|
| IFNα/β1 | Extracellular IFN-α/β1 |
| IFNγ | Extracellular IFN-γ |
| $R_{IFN\alpha/\beta1}$ | IFN-α/β1 receptor |
| $R_{IFN\gamma}$ | IFN-γ receptor |
| STAT1 | Unphosphorylated cytoplasmic STAT1 |
| STAT2 | Unphosphorylated cytoplasmic STAT2 |
| $STAT1_p$ | Phosphorylated cytoplasmic STAT1 |
| $STAT2_p$ | Phosphorylated cytoplasmic STAT2 |
| $(STAT1_p\|STAT1_p)$ | Cytoplasmic $STAT1_p\|STAT1_p$ complex |
| $(STAT1_p\|STAT2_p)$ | Cytoplasmic STAT1p\|STAT2p complex |
| IRF1 | Cytoplasmic active IRF1 protein |
| $IRF1_{inactive}$ | Cytoplasmic inactive IRF1 protein |
| $PTP_{active}$ | Cytoplasmic active PTP protein |
| $PTP_{inactive}$ | Cytoplasmic inactive PTP protein |
| $(PTP_{active}\|STAT1_p\|STAT1_p)$ | Cytoplasmic $PTP_{active}\|STAT1_p\|STAT1_p$ complex |
| $(PTP_{active}\|STAT1_p\|STAT2_p)$ | Cytoplasmic $PTP_{active}\|STAT1_p\|STAT2_p$ complex |
| $STAT1_n$ | Unphosphorylated nuclear STAT1 |
| $STAT2_n$ | Unphosphorylated nuclear STAT2 |
| $(STAT1_p\|STAT1_p)_n$ | Nuclear $STAT1_p\|STAT1_p$ complex |
| $(STAT1_p\|STAT2_p)_n$ | Nuclear STAT1p\|STAT2p complex |
| $IRF1_n$ | Nuclear active IRF1 protein |
| $IRF1_{inactive\_n}$ | Nuclear inactive IRF1 protein |
| $(STAT1\|IRF1)_n$ | Nuclear STAT1\|IRF1 complexes |
| $PTP_{active\_n}$ | Nuclear active PTP protein |
| $PTP_{inactive\_n}$ | Nuclear inactive PTP protein |
| $(PTP_{active}\|STAT1_p\|STAT1_p)$ | Nuclear $PTP_{active}\|STAT1_p\|STAT1_p$ complex |
| $(PTP_{active}\|STAT1_p\|STAT2_p)$ | Nuclear $PTP_{active}\|STAT1_p\|STAT2_p$ complex |
| $IRF1_t$ | IRF1 mRNA |
| $STAT1_t$ | STAT1 mRNA |
| $STAT2_t$ | STAT2 mRNA |

**Table S2. List of. model variables.** Variables refer to cytoplasmic content if no subscripts are present, subscript *n* denotes nuclear content, subscript *p* indicates a phosphorylated form, subscript *t* denotes mRNA transcripts, subscripts *active* and *inactive* are used to distinguish respective states of the molecules.

| No. | Parameter | Value | Source |
|---|---|---|---|
| 1 | $k_v$ | 1 | (Smieja *et. al*, 2008) |
| 2 | $v_{i1t}$ | $5.0 \cdot 10^{-6}$ [1/s] | (Smieja *et. al*, 2008) |
| 3 | $v_{s1t}$ | $2.3 \cdot 10^{-8}$ [1/s] | (Smieja *et. al*, 2008) |
| 4 | $v_{s2t}$ | $0.1 \cdot 10^{-5}$ [1/s] | (Smieja *et. al*, 2008) |
| 5 | $k_{transl}$ | 0.36 [1/s] | (Smieja *et. al*, 2008) |
| 6 | $k_{s1deg}$ | $1.28 \cdot 10^{-5}$ [1/s] | (Smieja *et. al*, 2008) |
| 7 | $k_{s1pdeg}$ | $1.28 \cdot 10^{-5}$ [1/s] | (Smieja *et. al*, 2008) |
| 8 | $k_{s2deg}$ | $1.28 \cdot 10^{-5}$ [1/s] | (Smieja *et. al*, 2008) |
| 9 | $k_{s2pdeg}$ | $1.28 \cdot 10^{-5}$ [1/s] | (Smieja *et. al*, 2008) |
| 10 | $k_{i1deg}$ | $0.58 \cdot 10^{-3}$ [1/s] | (Smieja *et. al*, 2008) |
| 11 | $k_{degs1t}$ | $9.71 \cdot 10^{-5}$ [1/s] | (Smieja *et. al*, 2008) |
| 12 | $k_{degs2t}$ | $9.71 \cdot 10^{-5}$ [1/s] | (Smieja *et. al*, 2008) |
| 13 | $k_{degi1t}$ | $6.41 \cdot 10^{-5}$ [1/s] | (Smieja *et. al*, 2008) |
| 14 | $k_{invs1s1}$ | $0.46 \cdot 10^{-3}$ [1/s] | (Smieja *et. al*, 2008) |
| 15 | $k_{invs1s1n}$ | $0.46 \cdot 10^{-3}$ [1/s] | (Smieja *et. al*, 2008) |
| 16 | $k_{invs1s2}$ | $0.46 \cdot 10^{-3}$ [1/s] | (Smieja *et. al*, 2008) |
| 17 | $k_{invs1s2n}$ | $0.46 \cdot 10^{-3}$ [1/s] | (Smieja *et. al*, 2008) |
| 18 | $k_{invs1i1}$ | $2.88 \cdot 10^{-3}$ [1/s] | (Smieja *et. al*, 2008) |
| 19 | $k_{s1s1pdeg}$ | $2.31 \cdot 10^{-5}$ [1/s] | (Smieja *et. al*, 2008) |
| 20 | $k_{s1s2pdeg}$ | $4.62 \cdot 10^{-5}$ [1/s] | (Smieja *et. al*, 2008) |
| 21 | $k_{s1i1deg}$ | $2.89 \cdot 10^{-5}$ [1/s] | (Smieja *et. al*, 2008) |
| 22 | $k_{s1s1}$ | 4 [1/(μM·s)] | (Smieja *et. al*, 2008) |
| 23 | $k_{s1s2}$ | 10 [1/(μM·s)] | (Smieja *et. al*, 2008) |
| 24 | $k_{s1i1}$ | 0.01 [1/(μM·s)] | (Smieja *et. al*, 2008) |
| 25 | $k_{inacti1}$ | $0.77 \cdot 10^{-3}$ [1/s] | (Smieja *et. al*, 2008) |
| 26 | $k_{s1tprod}$ | $1.67 \cdot 10^{-9}$ [μM/s] | (Smieja *et. al*, 2008) |
| 27 | $k_{s2tprod}$ | $1.53 \cdot 10^{-9}$ [μM/s] | (Smieja *et. al*, 2008) |
| 28 | $e_{s1}$ | 0.023 [1/s] | (Smieja *et. al*, 2008) |
| 29 | $i_{s1}$ | $0.23 \cdot 10^{-3}$ [1/s] | (Smieja *et. al*, 2008) |
| 30 | $e_{s2}$ | 0.023 [1/s] | (Smieja *et. al*, 2008) |
| 31 | $i_{s2}$ | $0.23 \cdot 10^{-3}$ [1/s] | (Smieja *et. al*, 2008) |
| 32 | $i_{s1s1}$ | 0.023 [1/s] | (Smieja *et. al*, 2008) |
| 33 | $i_{s1s2}$ | 0.023 [1/s] | (Smieja *et. al*, 2008) |
| 34 | $i_{i1}$ | 0.023 [1/s] | (Smieja *et. al*, 2008) |
| 35 | $e_{i1}$ | $2.3 \cdot 10^{-6}$ [1/s] | (Smieja *et. al*, 2008) |
| 36 | $e_{i1\_in}$ | $0.58 \cdot 10^{-2}$ [1/s] | (Smieja *et. al*, 2008) |
| 37 | $k_{s1phosRg}$ | 2.37 [1/s] | (Smieja *et. al*, 2008) |
| 38 | $k_{s1phosRab}$ | 1.66 [1/s] | fitted |
| 39 | $k_{s1dephc}$ | $0.58 \cdot 10^{-3}$ [1/s] | (Smieja *et. al*, 2008) |
| 40 | $k_{s2phosRab}$ | 1.18 [1/s] | (Smieja *et. al*, 2008) |
| 41 | $k_{s2dephc}$ | $0.58 \cdot 10^{-3}$ [1/s] | (Smieja *et. al*, 2008) |
| 42 | $k_{s1phos\_sat}$ | 10000 | (Smieja *et. al*, 2008) |
| 43 | $k_{s2phos\_sat}$ | 10000 | (Smieja *et. al*, 2008) |
| 44 | $R_{tot\_ab}$ | 0.1 [μM] | fitted |
| 45 | $k_{IFNab}$ | $1.6 \cdot 10^{-3}$ [1/s] | fitted |
| 46 | $k_{invIFNab}$ | $0.38 \cdot 10^{-3}$ [1/s] | fitted |
| 47 | $k_{IFNabdeg}$ | $0.14 \cdot 10^{-3}$ [1/s] | (Kumar *et. al*, 2003) |
| 48 | $R_{tot\_g}$ | 0.1 [μM] | fitted |

| | | | |
|---|---|---|---|
| 49 | $k_{IFNg}$ | $0.16 \cdot 10^{-2}$ [1/s] | fitted |
| 50 | $k_{invIFNg}$ | $9.63 \cdot 10^{-5}$ [1/s] | fitted |
| 51 | $k_{IFNgdeg}$ | $8.75 \cdot 10^{-5}$ [1/s] | (Londino *et. al*, 2018) |
| 52 | $R_{st}$ | $0.25 \cdot 10^{-2}$ [μM] | fitted |
| 53 | $k_{actPTP}$ | $2.8 \cdot 10^{-5}$ [1/s] | fitted |
| 54 | $k_{inactPTP}$ | 0 [1/s] | fitted |
| 55 | $k_{PTPss}$ | 0.35 [1/(μM·s)] | fitted |
| 56 | $k_{invPTPss}$ | $0.38 \cdot 10^{-2}$ [1/s] | fitted |
| 57 | $P_{TPinRa}$ | 7500 | fitted |
| 58 | $P_{TPinRg}$ | 1500 | fitted |
| 59 | $e_{PTP}$ | 0.019 [1/s] | fitted |
| 60 | $i_{PTP}$ | 0.058 [1/s] | fitted |
| ***Auxiliary equations parameters*** | | | |
| 61 | $k_1$ | $0.33 \cdot 10^{-3}$ [1/s] | (Smieja *et. al*, 2008) |
| 62 | $k_2$ | $0.65 \cdot 10^{-3}$ [1/s] | fitted |
| 63 | $k_3$ | $0.13 \cdot 10^{-2}$ [1/s] | fitted |
| 64 | $k_4$ | $0.49 \cdot 10^{-2}$ [1/s] | fitted |
| 65 | $k_5$ | $6.54 \cdot 10^{-5}$ [1/s] | fitted |

**Table S3. Model parameters.**

$$\frac{dIFN\alpha/\beta1}{dt} = -k_{IFNabdeg} \cdot IFN\alpha/\beta - k_{IFNab} \cdot IFN\alpha/\beta1 \cdot \left(R_{tot\_\alpha/\beta1} - R_{IFN\alpha/\beta1}\right) \cdot \frac{1}{1+k_{PTPinRab} \cdot PTP_{active}} \quad (S1.1)$$

$$\frac{dIFN\gamma}{dt} = -k_{IFNgdeg} \cdot IFN\gamma - k_{IFNg} \cdot IFN\gamma \cdot \left(R_{tot\_\gamma} - R_{IFN\gamma}\right) \cdot \frac{1}{1+k_{PTPinRg} \cdot PTP_{active}} \quad (S1.2)$$

$$\frac{dR_{IFN\alpha/\beta1}}{dt} = -k_{invIFNab} \cdot R_{IFN\alpha/\beta1} + k_{IFNab} \cdot IFN\alpha/\beta1 \cdot \left(R_{tot\_\alpha/\beta1} - R_{IFN\alpha/\beta1}\right) \cdot \frac{1}{1+k_{PTPinRab} \cdot PTP_{active}} \quad (S1.3)$$

$$\frac{dR_{IFN\gamma}}{dt} = -k_{invIFNg} \cdot R_{IFN\gamma} + k_{IFNg} \cdot IFN\gamma \cdot \left(R_{tot\_\gamma} - R_{IFN\gamma}\right) \cdot \frac{1}{1+k_{PTPinRg} \cdot PTP_{active}} \quad (S1.4)$$

$$\frac{dSTAT1}{dt} = -\left(k_{s1phosRab} \cdot \frac{R_{IFN\alpha/\beta1}}{R_{st}+R_{IFN\alpha/\beta1}} + k_{s1phosRg} \cdot \frac{R_{IFN\gamma}}{R_{st}+R_{IFN\gamma}}\right) \cdot \frac{STAT1}{1+k_{s1phossat} \cdot STAT1} + k_{s1dephc} \cdot STAT1_p + 2k_{invs1s1} \cdot (STAT1_p|STAT1_p) + k_{invs1s2} \cdot (STAT1_p|STAT2_p) + e_{s1} \cdot STAT1_n - i_{s1} \cdot STAT1 + k_{transl} \cdot STAT1_t - k_{s1deg} \cdot STAT1 + 2k_{invPTPss} \cdot (PTP_{active}|STAT1_p|STAT1_p) + k_{invPTPss} \cdot (PTP_{active}|STAT1_p|STAT2_p) \quad (S1.5)$$

$$\frac{dSTAT2}{dt} = -k_{s2phosRab} \cdot \frac{R_{IFN\alpha/\beta1}}{R_{st}+R_{IFN\alpha/\beta1}} \cdot \frac{STAT2}{1+k_{s2phossat} \cdot STAT2} + k_{s2dephc} \cdot STAT2_p + k_{invs1s2} \cdot (STAT1_p|STAT2_p) + e_{s2} \cdot STAT2_n - i_{s2} \cdot STAT2 + k_{transl} \cdot STAT2_t - k_{s2deg} \cdot STAT2 + k_{invPTPss} \cdot (PTP_{active}|STAT1_p|STAT2_p) \quad (S1.6)$$

$$\frac{dSTAT1_p}{dt} = \left(k_{s1phosRab} \cdot \frac{R_{IFN\alpha/\beta1}}{R_{st}+R_{IFN\alpha/\beta1}} + k_{s1phosRg} \cdot \frac{R_{IFN\gamma}}{R_{st}+R_{IFN\gamma}}\right) \cdot \frac{STAT1}{1+k_{s1phossat} \cdot STAT1} - k_{s1pdeg} \cdot STAT1_p - k_{s1dephc} \cdot STAT1_p - 2k_{s1s1} \cdot STAT1_p \cdot STAT1_p - k_{s1s2} \cdot STAT1_p \cdot STAT2_p \quad (S1.7)$$

$$\frac{dSTAT2_p}{dt} = k_{s2phosRab} \cdot \frac{R_{IFN\alpha/\beta1}}{R_{st}+R_{IFN\alpha/\beta1}} \cdot \frac{STAT2}{1+k_{s2phossat} \cdot STAT2} - k_{s2dephc} \cdot STAT2_p - k_{s1s2} \cdot STAT1_p \cdot STAT2_p - k_{s2pdeg} \cdot STAT2_p \quad (S1.8)$$

$$\frac{d(STAT1_p|STAT1_p)}{dt} = k_{s1s1} \cdot STAT1_p \cdot STAT1_p - k_{s1s1pdeg} \cdot (STAT1_p|STAT1_p) - i_{s1s1} \cdot (STAT1_p|STAT1_p) - k_{invs1s1} \cdot (STAT1_p|STAT1_p) - k_{PTPss} \cdot (STAT1_p|STAT1_p) \cdot PTP_{active} \quad (S1.9)$$

$$\frac{d(STAT1_p|STAT2_p)}{dt} = k_{s1s2} \cdot STAT1_p \cdot STAT2_p - k_{s1s2pdeg} \cdot (STAT1_p|STAT2_p) - i_{s1s2} \cdot (STAT1_p|STAT2_p) - k_{invs1s2} \cdot (STAT1_p|STAT2_p) - k_{PTPss} \cdot (STAT1_p|STAT2_p) \cdot PTP_{active} \quad (S1.10)$$

$$\frac{dIRF1}{dt} = k_{transl} \cdot IRF1_t - k_{i1deg} \cdot IRF1 - i_{i1} \cdot IRF1 + e_{i1} \cdot IRF1_n \quad (S1.11)$$

$$\frac{dIRF1_{inactive}}{dt} = -k_{i1deg} \cdot IRF1_{inactive} + e_{i1} \cdot IRF1_{inactive\_n} \quad (S1.12)$$

$$\frac{dPTP_{active}}{dt} = -i_{PTP} \cdot PTP_{active} + e_{PTP} \cdot PTP_{active\_n} + k_{actPTP} \cdot PTP_{inactive} \cdot x_{aux7} - k_{inactPTP} \cdot PTP_{active} - k_{PTPss} \cdot (STAT1_p|STAT1_p) \cdot PTP_{active} - k_{PTPss} \cdot (STAT1_p|STAT2_p) \cdot PTP_{active} + k_{invPTPss} \cdot (PTP_{active}|STAT1_p|STAT1_p) + k_{invPTPss} \cdot (PTP_{active}|STAT1_p|STAT2_p) \quad (S1.13)$$

$$\frac{dPTP_{inactive}}{dt} = -i_{PTP} \cdot PTP_{inactive} + e_{PTP} \cdot PTP_{inactive\_n} - k_{actPTP} \cdot PTP_{inactive} \cdot x_{aux7} + k_{inactPTP} \cdot PTP_{active} \quad (S1.14)$$

$$\frac{d(PTP_{active}|STAT1_p|STAT1_p)}{dt} = k_{PTPss} \cdot (STAT1_p|STAT1_p) \cdot PTP_{active} - k_{invPTPss} \cdot (PTP_{active}|STAT1_p|STAT1_p) \quad (S1.15)$$

$$\frac{d(PTP_{active}|STAT1_p|STAT2_p)}{dt} = k_{PTPss} \cdot (STAT1_p|STAT2_p) \cdot PTP_{active} - k_{invPTPss} \cdot (PTP_{active}|STAT1_p|STAT2_p) \quad (S1.16)$$

$$\frac{dSTAT1_n}{dt} = i_{s1} \cdot k_v \cdot STAT1 - e_{s1} \cdot k_v \cdot STAT1_n - k_{s1deg} \cdot STAT1_n + 2k_{invs1s1n} \cdot (STAT1_p|STAT1_p)_n + k_{invs1s2n} \cdot (STAT1_p|STAT2_p)_n + k_{invs1i1} \cdot (STAT1|IRF1)_n - k_{s1i1} \cdot IRF1_{active_n} \cdot STAT1_n + 2k_{invPTPss} \cdot (PTP_{active}|STAT1_p|STAT1_p)_n + k_{invPTPss} \cdot (PTP_{active}|STAT1_p|STAT2_p)_n \quad (S1.17)$$

$$\frac{dSTAT2_n}{dt} = i_{s2} \cdot k_v \cdot STAT2 - e_{s2} \cdot k_v \cdot STAT2_n - k_{s2deg} \cdot STAT2_n + k_{invs1s2n} \cdot (STAT1_p|STAT2_p)_n + k_{invPTPss} \cdot (PTP_{active}|STAT1_p|STAT2_p)_n \quad (S1.18)$$

$$\frac{d(STAT1_p|STAT1_p)_n}{dt} = -(k_{invs1s1} + k_{s1s1pdeg}) \cdot (STAT1_p|STAT1_p)_n + i_{s1s1} \cdot k_v \cdot (STAT1_p|STAT1_p) - k_{Ys1s1} \cdot (STAT1_p|STAT1_p)_n \cdot Y_{active_n} - k_{PTPss} \cdot (STAT1_p|STAT1_p)_n \cdot PTP_{active\_n} \quad (S1.19)$$

$$\frac{d(STAT1_p|STAT2_p)_n}{dt} = -(k_{invs1s2} + k_{s1s2pdeg}) \cdot (STAT1_p|STAT2_p)_n + i_{s1s2} \cdot k_v \cdot (STAT1_p|STAT2_p) - k_{PTPss} \cdot (STAT1_p|STAT2_p)_n \cdot PTP_{active\_n} \quad (S1.20)$$

$$\frac{dIRF1_n}{dt} = k_v \cdot i_{i1} \cdot IRF1 - k_{i1deg} IRF1_n - k_v \cdot e_{i1} \cdot IRF1_n - k_{inacti1} \cdot IRF1_n - k_{s1i1} \cdot IRF1_n \cdot STAT1_n + k_{invs1i1} \cdot (STAT1|IRF1)_n \quad (S1.21)$$

$$\frac{dIRF1_{inactive\_n}}{dt} = k_{inacti1} \cdot IRF1_n - k_{i1indeg} \cdot IRF1_{inactive\_n} - k_v \cdot e_{i1\_in} \cdot IRF1_{inactive\_n} \quad (S1.22)$$

$$\frac{d(STAT1|IRF1)_n}{dt} = k_{s1i1} \cdot IRF1_n \cdot STAT1_n - (k_{s1i1deg} + k_{invs1i1}) \cdot (STAT1|IRF1)_n \quad (S1.23)$$

$$\frac{dPTP_{active\_n}}{dt} = k_v \cdot i_{PTP} \cdot PTP_{active} - k_v \cdot e_{PTP} \cdot PTP_{active_n} - k_{PTPss} \cdot (STAT1_p|STAT1_p)_n \cdot PTP_{active\_n} - k_{PTPss} \cdot (STAT1_p|STAT2_p)_n \cdot PTP_{active\_n} + k_{invPTPss} \cdot (PTP_{active}|STAT1_p|STAT1_p)_n + k_{invPTPss} \cdot (PTP_{active}|STAT1_p|STAT2_p)_n \quad (S1.24)$$

| | |
|---|---|
| $\frac{\mathrm{dPTP}_{inactive\_n}}{\mathrm{dt}} = k_v \cdot \mathrm{i_{PTP}} \cdot \mathrm{PTP}_{inactive} - k_v \cdot \mathrm{e_{PTP}} \cdot \mathrm{PTP}_{inactive\_n}$ | (S1.25) |
| $\frac{\mathrm{d}(PTP_{active}\lvert\mathrm{STAT1_p}\rvert\mathrm{STAT1_p})_n}{\mathrm{dt}} = \mathrm{k_{PTPss}} \cdot (STAT1_p\lvert STAT1_p)_n \cdot PTP_{active} - \mathrm{k_{invPTPss}} \cdot (PTP_{active}\lvert\mathrm{STAT1_p}\rvert\mathrm{STAT1_p})_n$ | (S1.26) |
| $\frac{\mathrm{d}(PTP_{active}\lvert\mathrm{STAT1_p}\rvert\mathrm{STAT2_p})_n}{\mathrm{dt}} = \mathrm{k_{PTPss}} \cdot (STAT1_p\lvert STAT2_p)_n \cdot PTP_{active} - \mathrm{k_{invPTPss}} \cdot (PTP_{active}\lvert\mathrm{STAT1_p}\rvert\mathrm{STAT2_p})_n$ | (S1.27) |
| $\frac{\mathrm{dIRF11_t}}{\mathrm{dt}} = \mathrm{v_{i1t}} \cdot (\mathrm{STAT1_p}\lvert\mathrm{STAT1_p})_\mathrm{n} - \mathrm{k_{degi1t}} \cdot \mathrm{IRF1_t}$ | (S1.28) |
| $\frac{\mathrm{d}STAT1_t}{\mathrm{dt}} = k_{s1tprod} + v_{s1t} \cdot x_{aux4} - k_{degs1t} \cdot STAT1_t$ | (S1.29) |
| $\frac{\mathrm{d}STAT2_t}{\mathrm{dt}} = k_{s2tprod} - k_{degs1t} \cdot STAT2_t$ | (S1.30) |
| $\frac{\mathrm{dx_{aux1}}}{\mathrm{dt}} = k_1 \cdot \mathrm{IRF1_n} - k_1 \cdot \mathrm{x_{aux1}}$ | (S1.31) |
| $\frac{\mathrm{dx_{aux2}}}{\mathrm{dt}} = k_1 \cdot \mathrm{x_{aux1}} - k_1 \cdot \mathrm{x_{aux2}}$ | (S1.32) |
| $\frac{\mathrm{dx_{aux3}}}{\mathrm{dt}} = k_1 \cdot \mathrm{x_{aux2}} - k_1 \cdot \mathrm{x_{aux3}}$ | (S1.33) |
| $\frac{\mathrm{dx_{aux4}}}{\mathrm{dt}} = k_1 \cdot \mathrm{x_{aux3}} - k_1 \cdot \mathrm{x_{aux4}}$ | (S1.34) |
| $\frac{\mathrm{dx_{aux5}}}{\mathrm{dt}} = k_3 \cdot \frac{R_{\mathrm{IFN}\alpha/\beta1}}{R_{st}+R_{\mathrm{IFN}\alpha/\beta1}} + k_4 \cdot \frac{R_{\mathrm{IFN}\gamma}}{R_{st}+R_{\mathrm{IFN}\gamma}} - k_2 \cdot \mathrm{x_{aux5}}$ | (S1.35) |
| $\frac{\mathrm{dx_{aux6}}}{\mathrm{dt}} = k_5 \cdot \mathrm{x_{aux5}} - k_1 \cdot \mathrm{x_{aux6}}$ | (S1.36) |
| $\frac{\mathrm{dx_{aux7}}}{\mathrm{dt}} = k_5 \cdot \mathrm{x_{aux6}} - k_1 \cdot \mathrm{x_{aux7}}$ | (S1.37) |

**Table S4. Mathematical model of JAK-STAT signalling.** The transport of mRNA to the cytoplasm is assumed to be very fast in relation to other processes and therefore neglected in the model. Auxiliary variables $x_{auxi}$ have been introduced to represent multistage processes that lead to time delays in PTP activation and between transcription factor and corresponding transcript peaks. Equations in black colour correspond to the original model by (Smieja *et. al*, 2008), red colour denotes new components.

| Variable: | Initial condition [μM]: |
|---|---|
| IFNα/β1 | 0 |
| IFNγ | 0 |
| $R_{IFNα/β1}$ | 0 |
| $R_{IFNγ}$ | 0 |
| STAT1 | 0.66 |
| STAT2 | 0.44 |
| $STAT1_p$ | 0 |
| $STAT2_p$ | 0 |
| $(STAT1_p\|STAT1_p)$ | 0 |
| $(STAT1_p\|STAT2_p)$ | 0 |
| IRF1 | 0 |
| $IRF1_{inactive}$ | 0 |
| $PTP_{active}$ | *0* |
| $PTP_{inactive}$ | *0.01* |
| $(PTP_{active}\|STAT1_p\|STAT1_p)$ | *0* |
| $(PTP_{active}\|STAT1_p\|STAT2_p)$ | *0* |
| $STAT1_n$ | $0.66 \cdot 10^{-2}$ |
| $STAT2_n$ | $0.44 \cdot 10^{-2}$ |
| $(STAT1_p\|STAT1_p)_n$ | *0* |
| $(STAT1_p\|STAT2_p)_n$ | *0* |
| $IRF1_n$ | *0* |
| $IRF1_{inactive\_n}$ | *0* |
| $(STAT1\|IRF1)_n$ | *0* |
| $PTP_{active\_n}$ | *0* |
| $PTP_{inactive\_n}$ | *0.03* |
| $(PTP_{active}\|STAT1_p\|STAT1_p)_n$ | *0* |
| $(PTP_{active}\|STAT1_p\|STAT2_p)_n$ | *0* |
| $IRF1_t$ | *0* |
| $STAT1_t$ | $1.72 \cdot 10^{-5}$ |
| $STAT2_t$ | $1.58 \cdot 10^{-5}$ |

**Table S5. Model initial conditions.**
